# Evolution of (p)ppGpp-HPRT regulation through diversification of an allosteric oligomeric interaction

**DOI:** 10.1101/621474

**Authors:** Brent W. Anderson, Kuanqing Liu, Christine Wolak, Katarzyna Dubiel, Kenneth A. Satyshur, James L. Keck, Jue D. Wang

## Abstract

The signaling ligand (p)ppGpp binds diverse targets across bacteria, yet the mechanistic and evolutionary basis underlying these ligand-protein interactions remains poorly understood. Here we identify a novel (p)ppGpp binding motif in the enzyme HPRT, where (p)ppGpp shares identical binding residues for PRPP and nucleobase substrates to regulate purine homeostasis. Intriguingly, HPRTs across species share the conserved binding site yet strongly differ in ligand binding, from strong inhibition by basal (p)ppGpp levels to weak regulation at induced concentrations. Surprisingly, strong ligand binding requires an HPRT dimer-dimer interaction that allosterically opens the (p)ppGpp pocket. This dimer-dimer interaction is absent in the common ancestor but evolved to favor (p)ppGpp binding in the vast majority of bacteria. We propose that the evolutionary plasticity of oligomeric interfaces enables allosteric adjustment of ligand regulation, bypassing constraints of the ligand binding site. Since most ligands bind near protein-protein interfaces, this principle likely extends to other protein-ligand interactions.

## INTRODUCTION

Regulation of proteins by signaling ligands is a universal mechanism that has evolved for rapid adaptation to changing conditions (Chubukov et al., 2014; Traut, 2008). How proteins evolve to recognize ligands is under ongoing investigation (Najmanovich, 2017; Taute et al., 2014).

A key signaling ligand in bacteria is the nucleotide (p)ppGpp, which can be rapidly induced upon starvation to elicit the stringent response, a global alteration of transcription and metabolism (Cashel et al., 1996; Gourse et al., 2018; Liu et al., 2015a). (p)ppGpp directly binds and regulates diverse targets, including RNA polymerase, DNA primase, GTPases, various metabolic enzymes, and riboswitches (Corrigan et al., 2016; Gourse et al., 2018; Liu et al., 2015a; Sherlock et al., 2018; Wang et al., 2018; Zhang et al., 2018). (p)ppGpp binds these effectors at positions ranging from protein interfaces to active sites with diverse themes that have not been systematically elucidated. In addition, the spectra of (p)ppGpp targets differ among bacterial species. For example, in Proteobacteria (p)ppGpp binds to RNA polymerase, but most other bacteria lack a direct (p)ppGpp-RNA polymerase interaction and (p)ppGpp instead regulates guanylate kinase (Gourse et al., 2018). Finally, (p)ppGpp produced at basal levels can also have important protective roles (Gaca et al., 2013, 2015a; Kriel et al., 2012; Potrykus et al., 2011; Puszynska and O’Shea, 2017), but molecular targets and mechanisms of basal inhibition have not been delineated.

Here we reveal how specificity of (p)ppGpp for its target can be established by characterizing (p)ppGpp regulation of the enzyme hypoxanthine phosphoribosyltransferase (HPRT). HPRT was one of the earliest identified targets of (p)ppGpp whose regulation enables cellular homeostasis (Hochstadt-Ozer and Cashel, 1972; Kriel et al., 2012). We identify a novel class of (p)ppGpp binding motif at the enzyme’s active site, and demonstrate that HPRTs from diverse bacterial phyla are highly sensitive to (p)ppGpp. Mechanistic and evolutionary analyses reveal that regulation by basal levels of (p)ppGpp also requires an HPRT dimer-dimer interaction that allosterically positions a flexible loop to allow strong (p)ppGpp binding, and the few bacterial HPRTs lacking this dimer-dimer interaction are largely refractory to (p)ppGpp regulation. We propose a principle of “oligomeric allostery” where protein oligomerization affects conformation of the ligand binding site. This principle may be applicable to many other proteins with broad implications in evolutionary diversification of oligomeric structures.

## RESULTS

### (p)ppGpp regulation of HPRT is conserved across bacteria and beyond

HPRT is a purine salvage enzyme that converts purine bases and PRPP to precursors of the essential nucleotide GTP, and its activity was previously found to be inhibited by (p)ppGpp in several bacteria to regulate an important component of bacterial fitness: GTP homeostasis (Gaca et al., 2015b; Hochstadt-Ozer and Cashel, 1972; Kriel et al., 2012). It has been shown for the (p)ppGpp targets RNA polymerase and guanylate kinase that conservation of their binding to (p)ppGpp is limited within distinct bacterial phyla (Liu et al., 2015b; Ross et al., 2016). We therefore tested the conservation of HPRT regulation by conducting a broad biochemical survey of HPRT enzymes from free-living bacteria, human commensals, and pathogens. We revealed (p)ppGpp can bind and inhibit activities of HPRTs from every phylum of bacteria we tested, including Actinobacteria, Bacteroidetes, Firmicutes, Deinococcus-Thermus, and Proteobacteria, and even eukaryotic HPRTs (Figure 1). To do so, we purified 32 recombinant HPRTs and tested their enzymatic activities and found that nearly all were inhibited by ppGpp and pppGpp (Figure 1A-C and Table 1). We also quantified their binding affinity (K_d_) for pppGpp with <u>d</u>ifferential <u>ra</u>dial <u>c</u>apillary <u>a</u>ction of ligand <u>a</u>ssay (DRaCALA) (Figure 1D and 1F) (Roelofs et al., 2011), results of which were highly comparable with well-established quantitative methods such as isothermal titration calorimetry (Figure 1E and Figure 1 – figure supplement 1). Using this method, we found that most HPRTs bind pppGpp with K_d_ values ranging from ≈0.1 to ≈10 μM (Figure 1F and Table 1), in agreement with their inhibition by (p)ppGpp. Considering that (p)ppGpp is induced to millimolar concentrations during starvation and its uninduced levels are ≈10 – 20 μM during exponential growth in *B. subtilis* and *E. coli* (Cashel et al., 1996; Kriel et al., 2012), these results suggest significant inhibition of HPRT under physiological, basal levels of (p)ppGpp.

**Figure 1.**
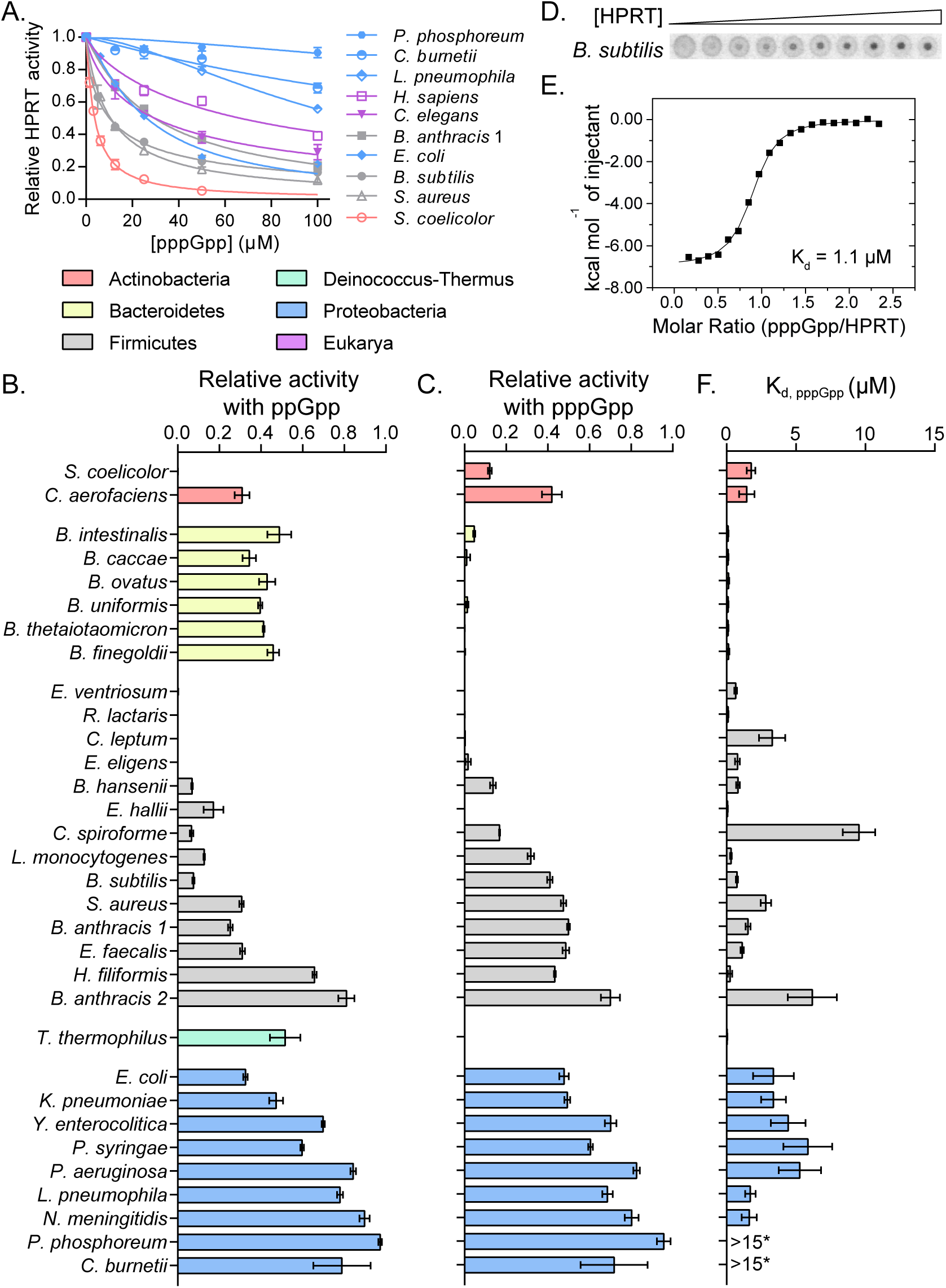
(p)ppGpp inhibition and binding to HPRT is conserved across species. **(A-C)** (p)ppGpp inhibits HPRTs across species. **A)** Relative activities of HPRTs with increasing concentrations of pppGpp. **B)** Relative activities of HPRTs with 25 μM ppGpp. **C)** Relative activities of HPRTs with 25 μM pppGpp. In A-C, error bars represent SEM of triplicates. **(D-F)** pppGpp binds HPRTs across species. **D)** Representative DRaCALA between *B. subtilis* HPRT in cell lysate serially diluted 1:2 and ^32^P-labeled pppGpp. The central signal is proportional to pppGpp – HPRT interaction. **E)** Binding isotherm from isothermal titration calorimetry between *B. anthracis* Hpt-1 and pppGpp. See Figure Supplement 1 for energy isotherm and parameters. **F)** The K_d_ between pppGpp and HPRTs obtained with DRaCALA from serially diluted cell lysates containing overexpressed HPRTs (see Materials and Methods). Error bars represent SEM derived from one binding curve. *P. phosphoreum* and *C. burnetii* HPRTs with * have affinities too weak to calculate but are estimated to be >15 μM (see Figure Supplement 1). See Table S1 for partial datasets for HPRTs from *M. tuberculosis*, *C. crescentus*, *S. meliloti*, *R. torques*, *S. mutans*, and *C. gilvus*.

**Table 1.** (p)ppGpp binding and inhibition of HPRT homologs.

| Organism | Relative activity<sup>d</sup> |  | K<sub>d</sub> (pppGpp)<br>(μM) | IC<sub>50</sub> (pppGpp)<br>(μM) |
| --- | --- | --- | --- | --- |
|  | 25 μM<br>ppGpp | 25 μM<br>pppGpp |  |  |
| **Actinobacteria** |  |  |  |  |
| *Streptomyces coelicolor* | -0.01 ± 0.01 | 0.12 ± 0.01 | 1.77 ± 0.31 | 3.8 ± 0.2 |
| *Collinsella aerofaciens* | 0.31 ± 0.04 | 0.42 ± 0.05 | 1.46 ± 0.56 |  |
| *Cellulomonas gilvus* | NA<sup>b</sup> | NA<sup>b</sup> | 0.6 ± 0.11 |  |
| *Mycobacterium tuberculosis* | NA<sup>a</sup> | NA<sup>a</sup> | NA<sup>c</sup> |  |
| **Bacteroidetes** |  |  |  |  |
| *Bacteroides intestinalis* | 0.49 ± 0.06 | 0.05 ± 0.01 | 0.11 ± 0.03 |  |
| *Bacteroides caccae* | 0.34 ± 0.03 | 0.01 ± 0.02 | 0.11 ± 0.02 |  |
| *Bacteroides ovatus* | 0.43 ± 0.04 | -0.03 ± 0.02 | 0.15 ± 0.04 |  |
| *Bacteroides uniformis* | 0.4 ± 0.01 | 0.01 ± 0.01 | 0.1 ± 0.04 |  |
| *Bacteroides thetaiotaomicron* | 0.41 ± 0 | 0 ± 0.01 | 0.11 ± 0.04 |  |
| *Bacteroides finegoldii* | 0.46 ± 0.03 | -0.01 ± 0.01 | 0.15 ± 0.06 |  |
| **Firmicutes** |  |  |  |  |
| *Eubacterium ventriosum* | -0.03 ± 0.03 | -0.07 ± 0.03 | 0.66 ± 0.1 |  |
| *Ruminococcus lactaris* | -0.01 ± 0 | 0 ± 0 | 0.11 ± 0.03 |  |
| *Ruminococcus torques* | NA<sup>b</sup> | NA<sup>b</sup> | 0.22 ± 0.07 |  |
| *Clostridium leptum* | -0.01 ± 0.01 | 0 ± 0 | 3.29 ± 0.94 |  |
| *Eubacterium eligens* | -0.03 ± 0.03 | 0.02 ± 0.01 | 0.79 ± 0.18 |  |
| *Blautia hansenii* | 0.07 ± 0 | 0.14 ± 0.01 | 0.83 ± 0.14 |  |
| *Eubacterium hallii* | 0.17 ± 0.05 | -0.06 ± 0.05 | 0.07 ± 0.02 |  |
| *Clostridium spiroforme* | 0.07 ± 0.01 | 0.17 ± 0 | 9.54 ± 1.17 |  |
| *Listeria monocytogenes* | 0.13 ± 0 | 0.32 ± 0.02 | 0.32 ± 0.06 |  |
| *Bacillus subtilis* | 0.08 ± 0.01 | 0.41 ± 0.01 | 0.75 ± 0.07 | 10.1 ± 1.3 |
| *Staphylococcus aureus* | 0.31 ± 0.01 | 0.48 ± 0.01 | 2.83 ± 0.38 | 10.6 ± 0.3 |
| *Streptococcus mutans* | NA<sup>b</sup> | NA<sup>b</sup> | 5.12 ± 1.45 |  |
| *Bacillus anthracis* 1 | 0.25 ± 0.01 | 0.5 ± 0.01 | 1.54 ± 0.18 | 29.5 ± 1.2 |
| *Enterococcus faecalis* | 0.31 ± 0.01 | 0.49 ± 0.02 | 1.12 ± 0.11 |  |
| *Holdemania filiformis* | 0.66 ± 0.01 | 0.43 ± 0 | 0.17 ± 0.11 |  |
| *Bacillus anthracis* 2 | 0.81 ± 0.04 | 0.7 ± 0.05 | 6.18 ± 1.77 |  |
| **Deinococcus-Thermus** |  |  |  |  |
| *Thermus thermophilus* | 0.52 ± 0.07 | -0.01 ± 0.01 | 0.04 ± 0.01 |  |
| **Proteobacteria** |  |  |  |  |
| *Caulobacter crescentus* | 0.33 ± 0.05 | 0.47 ± 0.07 | 0.24 ± 0.09 |  |
| *Escherichia coli* | 0.33 ± 0.01 | 0.48 ± 0.02 | 3.38 ± 1.47 | 26.2 ± 2.0 |
| *Klebsiella pneumoniae* | 0.47 ± 0.03 | 0.49 ± 0.01 | 3.38 ± 0.89 |  |
| *Sinorhizobium meliloti* | NA<sup>b</sup> | NA<sup>b</sup> | 3.08 ± 0.92 |  |
| *Yersinia enterocolitica* | 0.7 ± 0.01 | 0.7 ± 0.03 | 4.44 ± 1.25 |  |
| *Photobacterium phosphoreum* | 0.97 ± 0.01 | 0.96 ± 0.03 | NA<sup>c</sup> | >100 |
| *Pseudomonas syringae* | 0.6 ± 0.01 | 0.6 ± 0.01 | 5.85 ± 1.76 |  |
| *Pseudomonas aeruginosa* | 0.84 ± 0.01 | 0.83 ± 0.02 | 5.28 ± 1.52 |  |
| *Legionella pneumophila* | 0.78 ± 0.01 | 0.69 ± 0.03 | 1.72 ± 0.38 | >100 |
| *Coxiella burnetii* | 0.79 ± 0.14 | 0.72 ± 0.16 | NA<sup>c</sup> | >100 |
| *Neisseria meningitidis* | 0.9 ± 0.02 | 0.8 ± 0.03 | 1.64 ± 0.55 |  |
| **Eukarya** |  |  |  |  |
| *Saccharomyces cerevisiae* | 0.64 ± 0.02 | 0.96 ± 0.01 | 8.08 ± 8.58 |  |
| *Caenorhabditis elegans* | NA<sup>b</sup> | NA<sup>b</sup> | 12.77 ± 7.68 | 29.6 ± 2.8 |
| *Homo sapiens* | NA | NA | NA | 66.6 ± 5.4 |

± standard error of the mean
NA = no data obtained.
<sup>a</sup>Protein not purified
<sup>b</sup>Protein did not have activity
<sup>c</sup>Interaction with pppGpp too weak to estimate K<sub>d</sub>
<sup>d</sup>Relative to activity without ppGpp or pppGpp

Closer inspection revealed an additional quantitative difference in regulatory potency among these HPRTs that suggests a strong influence from the environmental niche occupied by the bacteria. HPRTs from human microbiota commensals, including all species from Bacteroidetes along with the Firmicutes *Collinsella aerofaciens*, *Eubacterium* spp., and *Ruminococcus* spp., manifested the tightest interactions with pppGpp (K_d_ ≈ 0.1 – 1 μM) (Figure 1F and Table 1). This suggests a relationship between the intestinal environment and regulation of this purine salvage enzyme by (p)ppGpp, which may serve to buffer the intracellular environment against fluctuating extracellular purine concentrations. Beyond microbiota, HPRTs from soil-dwelling bacteria (e.g., *Streptomyces coelicolor*, *B. subtilis*) and pathogens (e.g., *Bacillus anthracis, Staphylococcus aureus, E. coli*), occupying the phyla Actinobacteria, Firmicutes and Proteobacteria, were also inhibited by (p)ppGpp with K_d_ values below 10 μM (Figure 1B, 1C, and 1F). All eukaryotic HPRTs tested were inhibited by (p)ppGpp, although at reduced potency (Figure 1A and Table 1).

Interestingly, a few HPRTs were only weakly inhibited by (p)ppGpp, and we noticed that most of them come from intracellular pathogens that may not face purine fluctuations. These weakly regulated HPRTs all clustered in β- and γ-Proteobacteria (e.g., *Pseudomonas aeruginosa* and *Neisseria meningitidis*) (Figure 1B and 1C), with the exception of *Mycobacterium tuberculosis* (Figure 1 – figure supplement 1).

In addition to differences between HPRT homologs, we also noticed a strong, species-dependent difference between pppGpp and ppGpp inhibition. All HPRTs in Bacteroidetes were potently inhibited by pppGpp but only weakly by ppGpp (Figure 1B and 1C). In contrast, nearly all other HPRTs displayed stronger inhibition by ppGpp than pppGpp. This demonstrates that, although pppGpp and ppGpp are often regarded as similar, they can have marked differences for certain cellular targets.

### (p)ppGpp binds the conserved active site of HPRT and closely mimics substrate binding

To examine the molecular determinants underlying (p)ppGpp regulation of HPRT, we crystallized Hpt-1 from the pathogenic bacterium *B. anthracis* with and without ppGpp (Figure 2A and Table 2). For comparison, we also crystallized Hpt-1 with its two substrates, PRPP and the non-reactive guanine analog 9-deazaguanine (Figure 2B and Table 2) (Héroux et al., 2000). Apo HPRT diffracted to 2.06 Å resolution with four molecules in the asymmetric unit and was nearly identical to a deposited apo structure of *B. anthracis* Hpt-1 (PDB ID 3H83) (Figure 2 – figure supplement 1). The HPRT-substrates structure diffracted to 1.64 Å containing two molecules in the asymmetric unit (Figure 2 – figure supplement 1), and PRPP and 9-deazaguanine were coordinated by two Mg^2+^ ions with each monomer (Figure 2B). The HPRT-ppGpp complex diffracted to 2.1 Å resolution with two molecules in the asymmetric unit (Figure 2 – figure supplement 1), and each monomer contained one ppGpp coordinated with Mg^2+^, in agreement with near 1:1 stoichiometry measured via ITC (Figure 1 – figure supplement 1). These crystals formed in drops containing pppGpp, but there was insufficient density to completely model the 5′ γ-phosphate (Figure 2 – figure supplement 2). Although it is possible that the γ-phosphate was hydrolyzed during crystallization, the presence of waters and unassigned density around the 5′ phosphates allow for the possibility that the γ-phosphate is present but dynamic.

**Figure 2.**
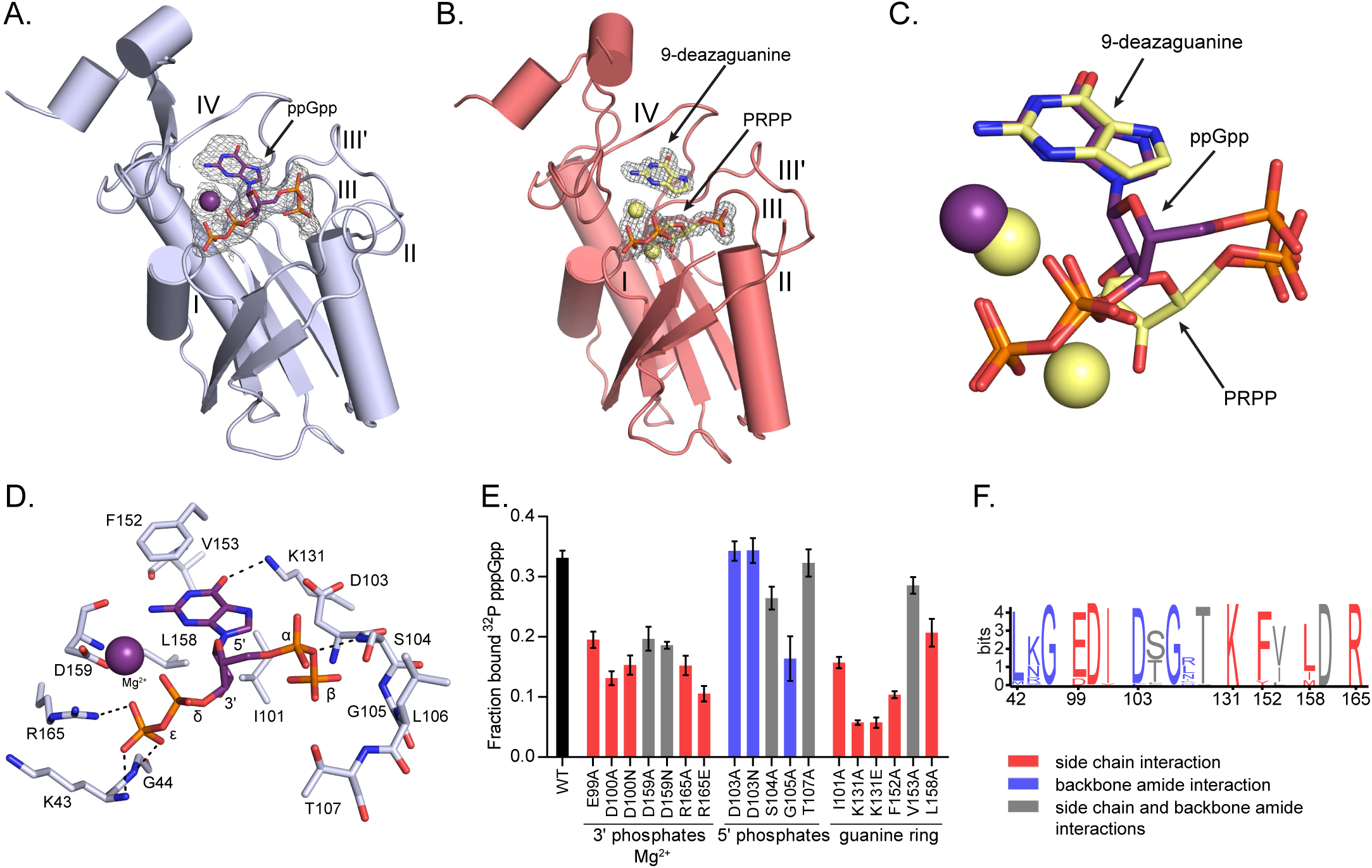
(p)ppGpp binds the HPRT active site and shares an almost identical binding pocket with substrates. **A)** *B. anthracis* Hpt-1 crystallized with ppGpp. Five loops (I, II, III, III′, IV) form the HPRT active site (Sinha and Smith, 2001). ppGpp and Mg^2+^ shown with omit electron density contoured at 2σ. Figure Supplement 2 shows omit electron density for the two ppGpp molecules crystallized in the asymmetric unit. **B)** *B. anthracis* Hpt-1 crystallized with 9-deazaguanine, PRPP, and two Mg^2+^. Ligands shown with omit electron density contoured at 2.5σ. Residues Tyr70 – Ser76 in loop II were not resolved. See Figure Supplement 1 for asymmetric units of Hpt-1 crystallized with ppGpp, substrates, and sulfates. **C)** Overlay of substrates (PRPP and 9-deazaguanine; yellow) and inhibitor (ppGpp; purple) bound to HPRT. Spheres represent Mg^2+^ and are colored according to their coordinating ligand. See Figure Supplement 4 for binding pocket comparison. **D)** The ppGpp binding pocket on HPRT. Black dotted lines indicate select hydrogen bonds. The peptide backbone is shown for residues where the interactions are relevant. See Figure Supplements 3 and 5 for complete interaction maps. **E)** DRaCALA of *B. subtilis* HPRT variants binding to ^32^P-labeled pppGpp, determined with cell lysates containing overexpressed HPRT variants. Disruption of side chain interactions weakened binding. For E and F, the residues are colored according to their interaction with ppGpp. Red = side chain interaction, blue = backbone amide interaction, and gray = both side chain and backbone interactions. Error bars represent SEM of three replicates. **F)** Sequence frequency logo of the (p)ppGpp binding pocket from 99 bacterial HPRTs. Logo created using WebLogo from UC-Berkeley. See Figure Supplement 6 for binding residues from select bacterial and eukaryotic HPRTs.

**Table 2.** X-ray data collection and structure determination statistics.

| **Data collection** |  |  |  |
| --- | --- | --- | --- |
| Structure | HPRT with sulfates<br>(PDB 6D9Q) | HPRT with substrates<br>(PDB 6D9R) | HPRT with ppGpp<br>(PDB 6D9S) |
| Wavelength | 0.9786 | 0.97872 | 0.9786 |
| Resolution range<br>(highest resolution<br>bin) (Å) | 41.3 – 2.056 (2.13 –<br>2.056) | 50 – 1.64 (1.67 –<br>1.64) | 40.2 – 2.105 (2.181 –<br>2.105) |
| Space group | P 3<sub>1</sub> 2 1 | P 3<sub>2</sub> 2 1 | P 3<sub>1</sub> 2 1 |
| Unit cell<br>a, b, c (Å) | 82.597, 82.597,<br>242.416 | 113.758, 113.758,<br>56.731 | 82.61, 82.61, 174.92 |
| $\alpha, \beta, \gamma$ (°) | 90, 90, 120 | 90, 90, 120 | 90, 90, 120 |
| Completeness (%) | 99.16 (95.52) | 99.8 (99.0) | 98.92 (94.51) |
| Unique reflections | 60264 | 51750 | 40413 |
| Redundancy | 10.8 (6.4) | 21.5 (14.4) | 8.1 (6.2) |
| $I/\sigma I$ | 24.8 (1.74) | 31.09 (2.93) | 16.41 (1.22) |
| R<sub>meas</sub> | 0.124 | 0.137 (1.376) | 0.164 |
| R<sub>pim</sub> | 0.037 | 0.029 (0.35) | 0.06 |
| CC 1/2 | (0.791) | (0.81) | (0.724) |
| **Refinement** |  |  |  |
| Resolution range<br>(highest resolution<br>bin) (Å) | 41.3 – 2.056 (2.107 –<br>2.056) | 37.19 – 1.640 (1.699<br>– 1.64) | 40.2 – 2.105 (2.158 –<br>2.105) |
| R<sub>work</sub>/R<sub>free</sub><sup>a</sup> (%) | 18.24 / 21.65 | 16.53 / 19.57 | 20.17 / 24.4 |
| r.m.s.<sup>b</sup> deviations |  |  |  |
| Bonds (Å) | 0.004 | 0.01 | 0.004 |
| Angles (Å) | 1.03 | 1.41 | 1.03 |
| Ramachandran<br>statistics (%) |  |  |  |
| Favored | 97.90 | 98.52 | 96.63 |
| Allowed | 2.10 | 1.48 | 3.37 |
| Disallowed | 0.00 | 0.00 | 0.00 |
| Rotamer outliers (%) | 0.78 | 0.30 | 0.63 |
| No. atoms |  |  |  |
| Macromolecules | 5742 | 2881 | 2837 |
| Ligands | 46 | 161 | 116 |
| Solvent | 440 | 317 | 118 |
| B factor (Å<sup>2</sup>) |  |  |  |
| Macromolecules | 50.63 | 24.06 | 76.03 |
| Ligands | 68.83 | 37.76 | 161.17 |
| Solvent | 52.15 | 35.53 | 70.22 |

<sup>a</sup>R<sub>work</sub>/R<sub>free</sub> = $\sum |F_{\text{obs}}| - |F_{\text{calc}}| / |F_{\text{obs}}|$ , where the working and free R factors are calculated by using the working and free reflection sets, respectively. The free R reflections were held aside throughout refinement. <sup>b</sup>Root mean square. See Source Data 1-3 for PDB validation reports.

Comparison between the HPRT-ppGpp and HPRT-substrates structures revealed that ppGpp binds the HPRT active site and closely mimics the conformation of the two substrates (Figure 2A-B and Figure 2 – figure supplement 3). The purine ring of ppGpp overlaps with the purine base substrate, and the two phosphate arms of ppGpp and PRPP both spread across the active site between loop I and loop III (Figure 2C and Figure 2 – figure supplement 4). In ppGpp-protein interactions, the phosphates of ppGpp are either elongated or compacted in a ring-like conformation (Steinchen and Bange, 2016). With the phosphates of ppGpp spread across the binding pocket, the HPRT-ppGpp interaction represents an elongated conformation (Figure 2A).

The (p)ppGpp binding site in HPRT manifests a novel (p)ppGpp-protein interaction. The 5′ phosphates and ribose of ppGpp interact with loop III (EDIIDSGLT), a well-characterized PRPP binding motif (Sinha and Smith, 2001), through side chain interactions with Glu99 and Asp100 and backbone amide interactions with Asp103 – Thr107 (Figure 2D and Figure 2 – figure supplement 5). The 3′ phosphates of ppGpp are coordinated by backbone amides of loop I, the side chain of Arg165, and the Mg^2+^ ion (Figure 2D and Figure 2 – figure supplement 5). The guanine ring of ppGpp is surrounded by a hydrophobic cleft formed by Ile101, Phe152, and Leu158, and Lys131 hydrogen bonds the guanine’s exocyclic oxygen (Figure 2D and Figure 2 – figure supplement 5). We validated the (p)ppGpp binding residues by mutation analyses which showed that altering residues with side chain interactions with (p)ppGpp greatly weakened pppGpp binding (Figure 2E). In sum, this binding pocket illustrates a novel (p)ppGpp binding motif. Since many proteins bind PRPP (Hove-Jensen et al., 2017), this (p)ppGpp motif may represent a new class of (p)ppGpp targets.

A frequency logo of the binding pocket from 99 bacterial HPRTs shows that most (p)ppGpp-interacting residues are highly conserved across bacteria (Figure 2F and Figure 2 – figure supplement 6), and nearly all binding residues are also conserved in the eukaryotic HPRTs (Figure 2 – figure supplement 6). The strong conservation across species is not surprising given the close overlap between (p)ppGpp and substrates. Since all residues involved in binding ppGpp are also involved in binding substrates (Figure 2 – figure supplement 3 and 4), altering the site to affect inhibitor binding would also impact enzyme activity.

The nearly identical recognition of (p)ppGpp and substrates, along with the conservation of the active site, raised the following question: how can some HPRTs be strongly inhibited by basal levels of (p)ppGpp and other HPRTs be almost refractory to (p)ppGpp control despite sharing a conserved binding pocket?

### (p)ppGpp prevents PRPP-induced dissociation of HPRT dimer-of-dimers

We noticed a significant difference between the ppGpp- and substrates-bound tertiary structures in the conformation of a flexible loop. This loop, also called loop II, is common to all HPRTs and covers the active site during catalysis (Shi et al., 1999). In the inhibited HPRT-ppGpp complex, one side of loop II faces the active site while the other side is a critical part of the interface between two HPRT dimers forming a tetramer (Figure 3A). In the substrates-bound state, loop II is instead shifted ~4 Å toward the active site from its position in the dimer-dimer interface (Figure 3A). Loop II also pulls its flanking dimer interface components β3 and α3 toward the active site (Figure 3A), abolishing the dimer-dimer interaction (Figure 3B). Using size-exclusion chromatography, we confirmed that apo *B. subtilis* HPRT is a tetramer. When PRPP, the first substrate to bind HPRT (Yuan et al., 1992), is added at a high concentration (500 μM) in the mobile phase, HPRT tetramers dissociate to dimers upon substrate binding (Figure 3C and Figure 3 – figure supplement 1 and 2).

**Figure 3.**
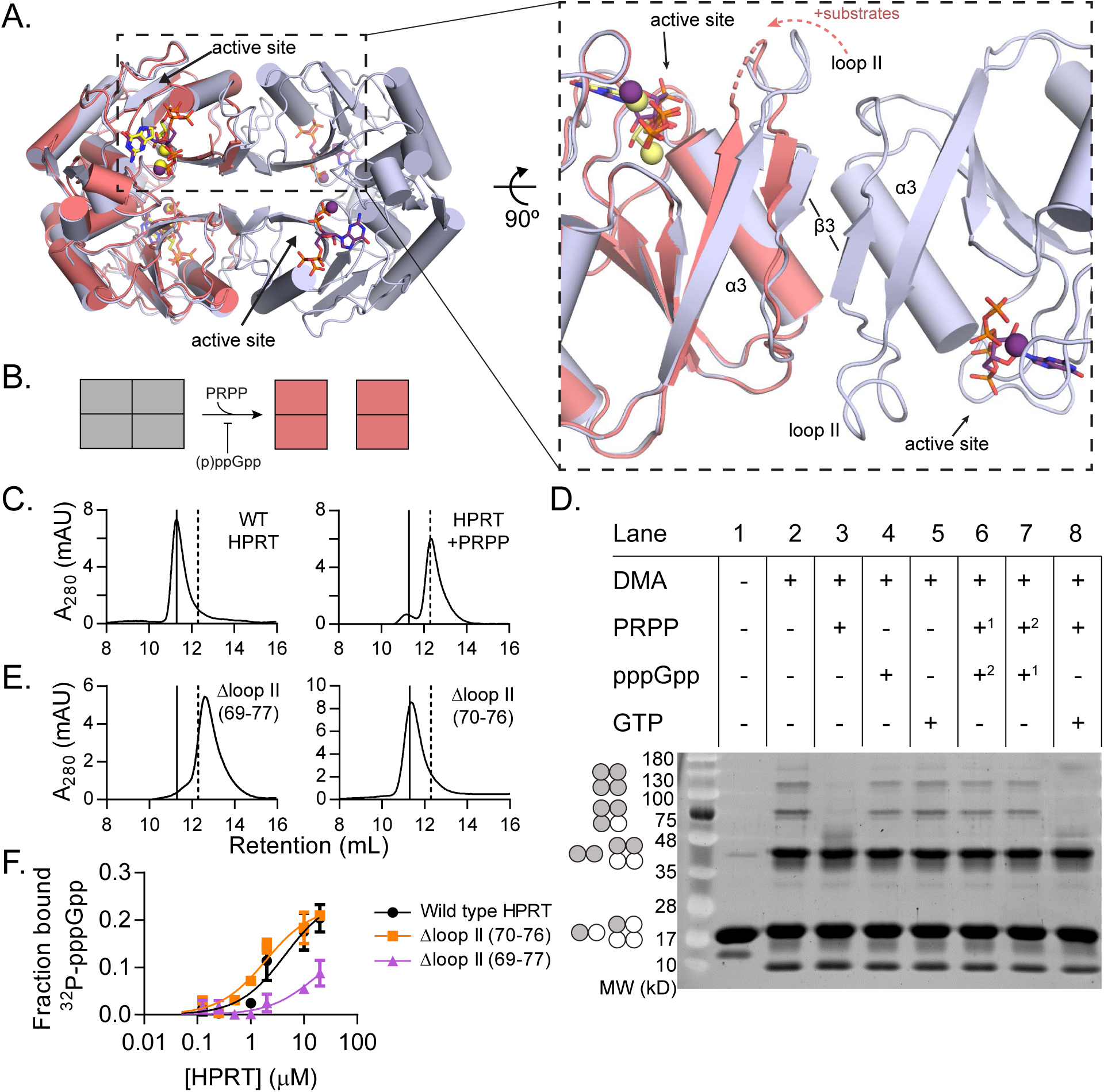
(p)ppGpp counteracts substrate-induced HPRT dimerization and HPRT tetramerization potentiates (p)ppGpp binding. **A)** Overlay of HPRT tetramer crystallized with ppGpp (silver) and HPRT dimer crystallized with substrates (salmon). ppGpp is purple and the substrates are yellow. Inset, View of the dimer-dimer interface within an HPRT tetramer. Secondary structure components at the interface are labeled β3, α3, and loop II. **B)** Schematic of tetramer to dimer transition dependent on PRPP and (p)ppGpp. **C)** Size-exclusion chromatographs of *B. subtilis* HPRT without ligand (left) and with PRPP (right). PRPP addition shifts the oligomeric state from tetramer to dimer. In C and E, solid line shows *B. subtilis* HPRT tetramer peak and dotted line shows dimer peak. PRPP must be in mobile phase to cause the shift (see Figure Supplement 2). For molecular weight standards, see Figure Supplement 3. *B. anthracis* Hpt-1 is also a dimer with PRPP (see Figure Supplement 1). **D)** *B. subtilis* HPRT crosslinked with dimethyl adipimidate (DMA). Crosslinked multimers were separated on SDS-PAGE. Predicted multimers are shown on the left. Shaded circles represent predicted to be crosslinked monomers, and white circles represent incomplete crosslinking. ^1, 2^ indicates order of incubation. **E)** Size-exclusion chromatography of *B. subtilis* HPRT with a partial loop II deletion (Δ70-76) or a complete loop II deletion (Δ69-77). The complete deletion results in dimerization of the protein. **F)** Binding curves between ^32^P-labeled pppGpp and *B. subtilis* HPRT variants obtained with DRaCALA. The tetrameric Δloop II (70-76) variant (orange squares) binds as well as wild type HPRT. The dimeric Δloop II (69-77) variant (purple triangles) displays weaker binding. Error bars represent SEM of three replicates.

We next interrogated the effect of (p)ppGpp on the oligomeric state of HPRT using the protein crosslinker dimethyl adipimidate (DMA). Crosslinked apo HPRT was resolved by SDS-PAGE as bands corresponding to monomers, dimers, trimers, or tetramers (Figure 3D, lane 2). All but the tetramer is likely formed by incomplete crosslinking, since dynamic light scattering showed apo *B. anthracis* Hpt-1 to be a homogeneous population with a hydrodynamic radius (R_H_) consistent with a tetramer in solution (Table 3). HPRT with pppGpp remained a tetramer according to both crosslinking and dynamic light scattering experiments (Figure 3D and Table 3). Incubation of HPRT with PRPP resulted in loss of trimer and tetramer bands in SDS-PAGE (Figure 3D, lane 3) and a corresponding shift in tetramer to homogeneous dimer with dynamic light scattering (Table 3), confirming that PRPP-bound HPRT is a dimer.

**Table 3.**
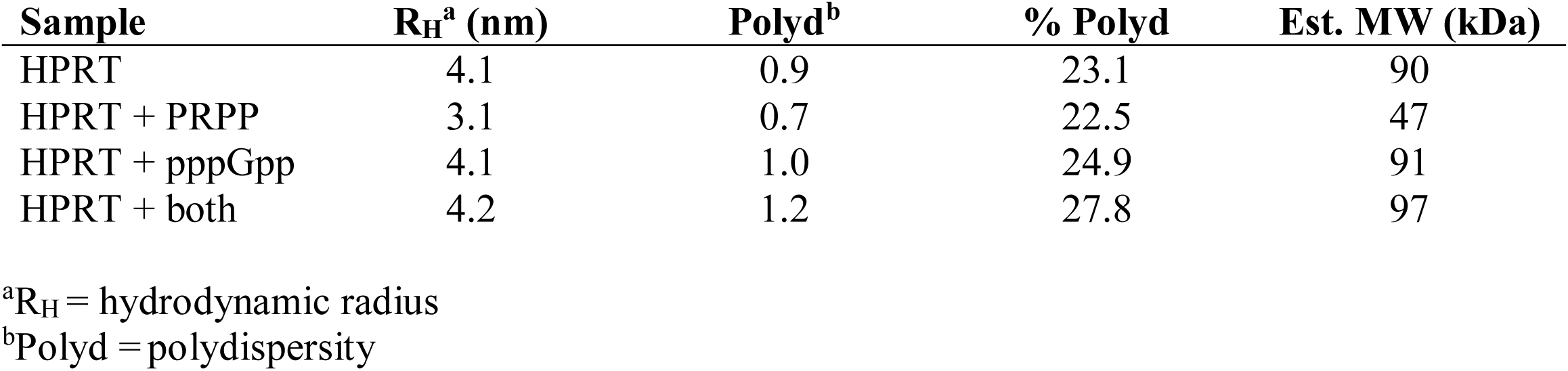
Dynamic light scattering of *B. anthracis* Hpt-1 with pppGpp and PRPP.

| **Sample** | **R<sub>H</sub><sup>a</sup> (nm)** | **Polyd<sup>b</sup>** | **% Polyd** | **Est. MW (kDa)** |
| --- | --- | --- | --- | --- |
| HPRT | 4.1 | 0.9 | 23.1 | 90 |
| HPRT + PRPP | 3.1 | 0.7 | 22.5 | 47 |
| HPRT + pppGpp | 4.1 | 1.0 | 24.9 | 91 |
| HPRT + both | 4.2 | 1.2 | 27.8 | 97 |

<sup>a</sup>R<sub>H</sub> = hydrodynamic radius
<sup>b</sup>Polyd = polydispersity

Importantly, with both PRPP and pppGpp present, HPRT was tetrameric (Figure 3D, lanes 6 and 7 and Table 3), indicating that pppGpp prevents PRPP-induced dimer-dimer dissociation. GTP competition with PRPP did not recapitulate pppGpp blockage of PRPP-induced dimer-dimer dissociation (Figure 3D, lane 8). Thus pppGpp appears to selectively stabilize HPRT tetramers against PRPP-induced dissociation. Taken together, these data reveal that HPRT binds PRPP as catalytically active dimers, whereas (p)ppGpp maintains HPRT as inactive tetramers, thus preventing the formation of active HPRTs.

### Dimer-dimer interaction allosterically positions loop II for potentiated (p)ppGpp binding

Given HPRT’s shift in oligomeric state with substrates and counteraction by (p)ppGpp, we reasoned that it could be an allosteric enzyme (Traut, 2008). Therefore, we examined whether there is cooperativity for substrate or inhibitor binding, a common feature of allosteric enzymes. However, ITC experiments with pppGpp did not suggest cooperative binding sites (Figure 1 – figure supplement 1). In addition, HPRT does not show positive cooperativity with respect to either substrate (Guddat et al., 2002; Patta et al., 2015). Therefore, HPRT is not an allosteric enzyme with respect to both substrates and inhibitor.

Despite not exhibiting features of an allosteric enzyme, we found that, unexpectedly, the oligomeric interaction is critical for HPRT’s regulation by (p)ppGpp. We disrupted the dimer-dimer interface of *B. subtilis* HPRT by constructing two loop II deletion variants of different lengths that resulted in different apo oligomeric states (Figure 3E). While the tetrameric Δloop II (70-76) bound pppGpp as tightly as wild type HPRT, the dimeric Δloop II (69-77) displayed strongly ablated binding to pppGpp (K_d_ too weak to estimate; Figure 3F).

Since an engineered dimeric HPRT variant has weakened binding to (p)ppGpp, we next examined the oligomeric state of naturally occurring HPRT homologs with weakened inhibition by (p)ppGpp (Figure 1). Strikingly, in nearly all cases the (p)ppGpp-insensitive HPRTs were also constitutive dimers (Figure 4A). Our results suggest that tetrameric HPRT, but not dimeric HPRT, allows (p)ppGpp at basal levels to bind to and inhibit its activity.

**Figure 4.**
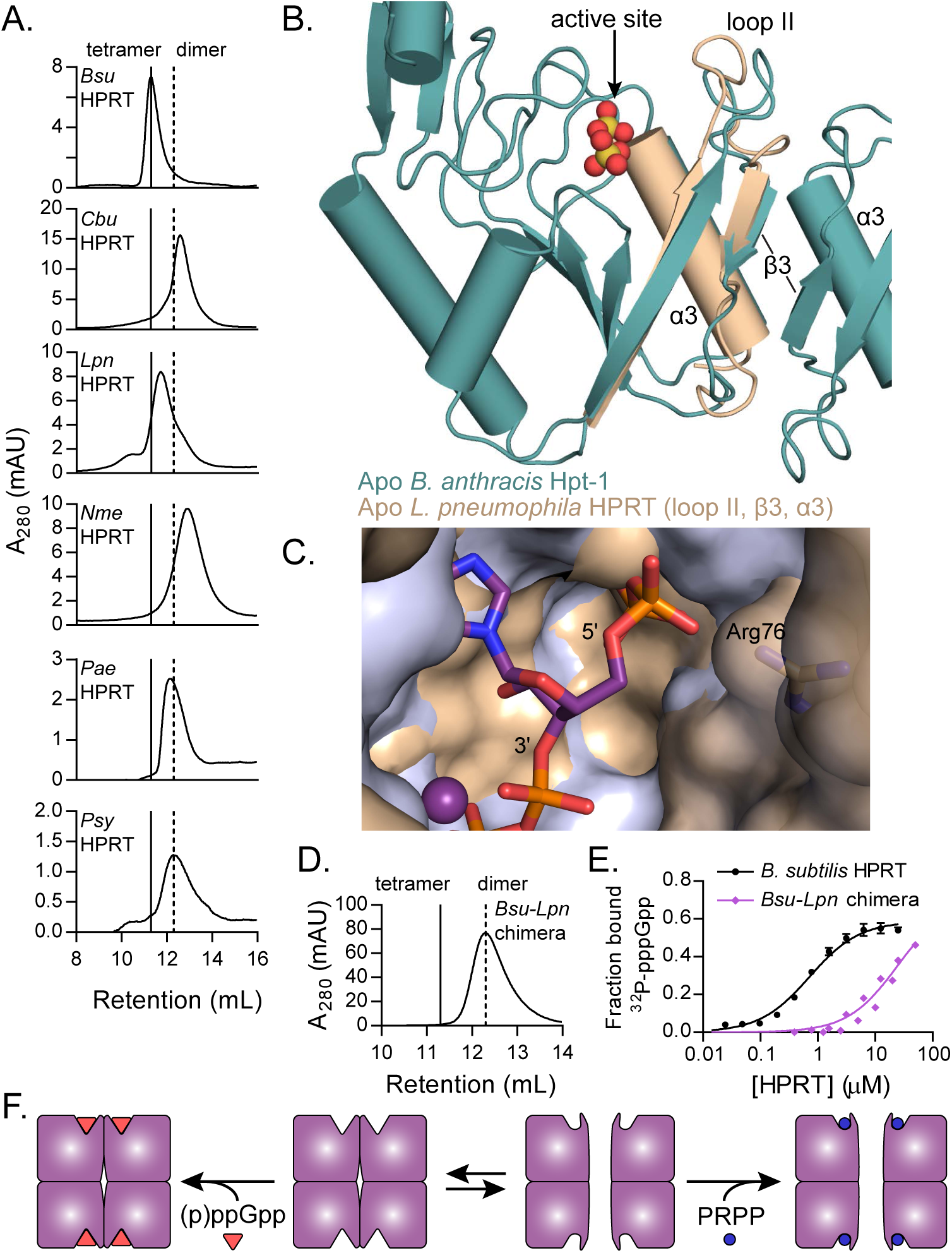
Dimer-dimer interaction holds loop II away from the (p)ppGpp binding pocket. **A)** Size-exclusion chromatographs show that *C. burnetii*, *L. pneumophila*, *N. meningitidis*, *P. aeruginosa*, and *P. syringae* HPRTs are non-tetrameric without ligands. Vertical lines represent *B. subtilis* HPRT tetramer and dimer peaks. **B)** Overlay of apo *B. anthracis* Hpt-1 (teal) and the *L. pneumophila* HPRT interface (wheat; PDB ID 5ESW). In *L. pneumophila* HPRT, loop II and β3 are further from the dimer-dimer interface and closer to the active site. See Figure Supplement 1 for overlay with substrates-bound HPRT. **C)** Overlay of *B. anthracis* Hpt-1 – ppGpp (silver; ppGpp in purple) and *L. pneumophila* HPRT (wheat; PDB ID 5ESX) shows that the 5′ phosphate binding pocket is compressed by the conformation of loop II in the dimeric HPRT. Arg76 (side chain shown as sticks) in *L. pneumophila* HPRT forms part of the compressed pocket. Asp109 in *L. pneumophila* HPRT is hidden due to poor electron density. **D)** Size-exclusion chromatograph of the dimeric *Bsu*-*Lpn* chimera. The chimera has 21 residues at the *B. subtilis* HPRT interface replaced with *L. pneumophila* HPRT residues (see Figure Supplement 2). **E)** DRaCALA shows that the *Bsu*-*Lpn* chimera binds ^32^P-labeled pppGpp more weakly. Error bars represent SEM of three replicates. *Bsu*-*Lpn* chimera performed in duplicate. Figure Supplement 3 shows that stability of dimeric HPRTs is not compromised. **F)** Model for how oligomerization affects (p)ppGpp binding. In HPRT’s tetrameric state, the dimer-dimer interface sequesters loop II at the interface, making the binding pocket conducive to (p)ppGpp binding. In its dimeric state, loop II is not sequestered at the interface, interfering with (p)ppGpp binding but not interaction with the substrate PRPP.

The structural basis of how the dimer-dimer interaction promotes (p)ppGpp binding in tetrameric HPRTs can be seen from comparative structures of apo *B. anthracis* Hpt-1 and apo *L. pneumophila* HPRT (PDB ID 5ESW) (Zhang et al., 2016). Loop II in dimeric *L. pneumophila* HPRT is positioned closer to the active site (Figure 4B), mimicking substrates-bound HPRT even in the absence of substrates (Figure 4 – figure supplement 1), and compresses the cavity surrounding the 5′ phosphates of ppGpp (Figure 4C). This conformation does not fully accommodate (p)ppGpp, but it should accommodate PRPP since its 5′ monophosphate fits in the cavity compressed by loop II (Figure 4 – figure supplement 1). In contrast, in tetrameric HPRTs, loop II is pulled away from the active site by the dimer-dimer interaction and is positioned for optimal (p)ppGpp binding (Figure 4F). While there are many examples of ligands affecting oligomerization (Traut, 1994b), here we have shown that for HPRT, its oligomeric state can instead affect ligand binding. Because the ligand binds to a non-interface pocket, the oligomeric state affects the ligand binding through allosteric interaction, which we will refer as “oligomeric allostery”.

To identify the determinant for the oligomeric state of naturally occurring HPRTs, we constructed a chimera of *B. subtilis* HPRT with 21 dimer-dimer interface residues replaced with their corresponding *L. pneumophila* HPRT residues (Figure 4 – figure supplement 2). The chimera is a stable dimer (Figure 4D and Figure 4 – figure supplement 3), indicating that the determinants for oligomerization lie in the dimer-dimer interface residues. The chimera also had a >20-fold lower affinity for pppGpp than the tetramer (K_d_ ≈ 24 μM versus ≈ 1 μM) (Figure 4E), suggesting that there may be an evolved linkage between the dimer-dimer interface of HPRT tetramers and (p)ppGpp binding.

### A dimer-dimer interface motif coevolved with strong (p)ppGpp binding across species

The relationship between HPRT oligomeric state and sensitivity to (p)ppGpp prompted us to examine whether HPRT oligomerization has coevolved with (p)ppGpp regulation. To test this, we turned to ancestral protein sequence reconstruction to infer the evolution of HPRT (Hochberg and Thornton, 2017). With a phylogenetic tree derived from an alignment of 141 bacterial HPRTs and *S. cerevisiae* HPRT as an outgroup, we used maximum likelihood to infer the most likely ancestral HPRT protein sequences based on the phylogeny of the extant HPRT sequences (Jones et al., 1992) (Figure 5A). From the first ancestral HPRT (Anc1), the phylogenetic tree bifurcated into two broad lineages (Figure 5A). One lineage contained the known dimeric HPRTs with weakened (p)ppGpp inhibition (Hug et al., 2016) (Figure 5A). The second lineage contained the vast majority of HPRTs, including (p)ppGpp-sensitive HPRTs in the Bacteroidetes, Firmicutes, and γ-proteobacteria. Notably, Anc1 HPRT was likely dimeric, since the interface of Anc1 shares three residues conserved only in the dimeric lineage (Trp82, Pro86, and Ala110) (Figure 5 – figure supplement 1). Varying these residues in *B. subtilis* HPRT produces non-tetrameric variants (Figure 5 – figure supplement 2).

**Figure 5.**
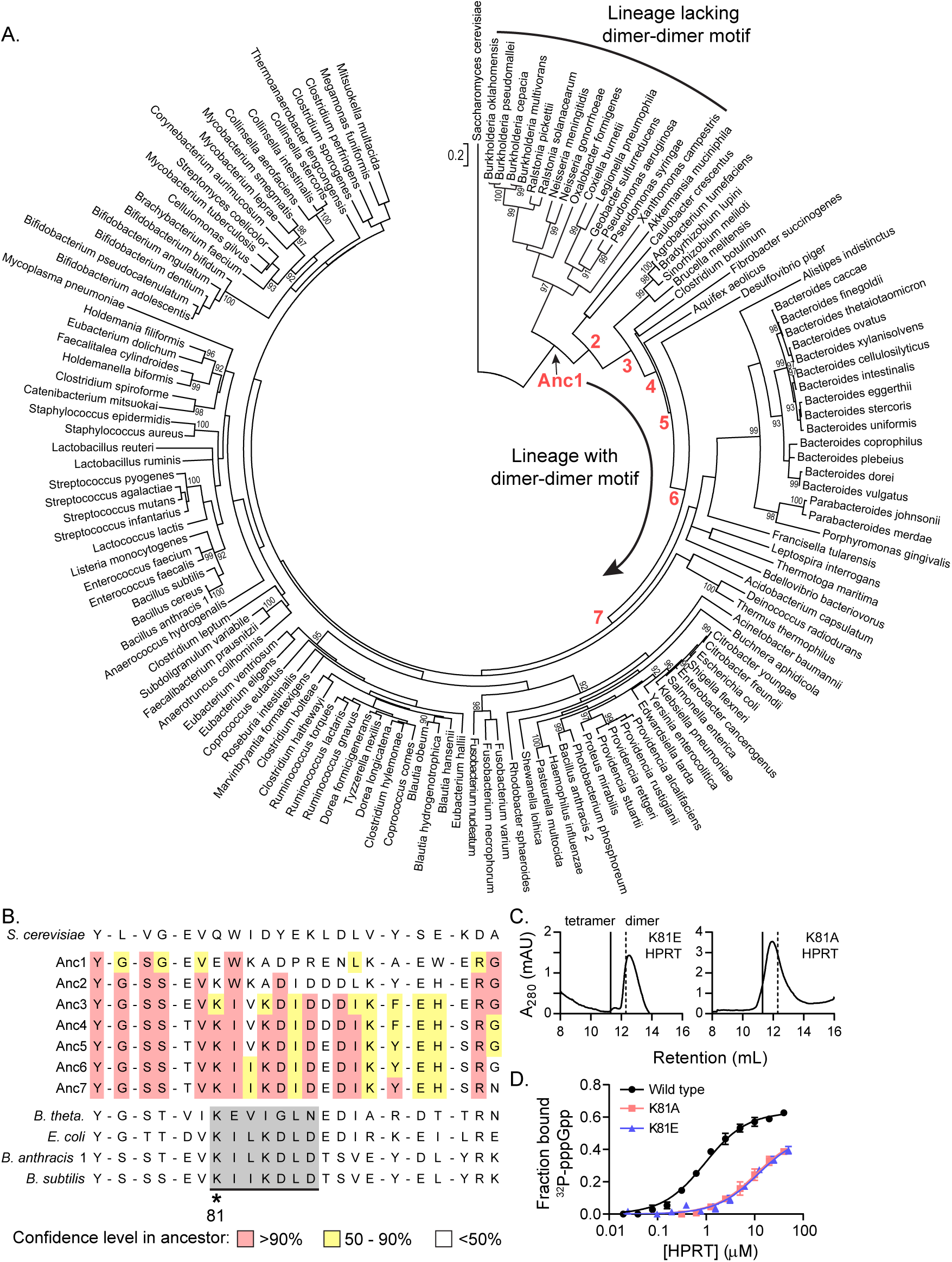
Coevolution of HPRT dimer-dimer interaction and (p)ppGpp regulation. **A)** A maximum likelihood phylogenetic tree of 141 bacterial HPRT amino acid sequences rooted on *Saccharomyces cerevisiae* HPRT. Bootstrap values greater than 90% from 1000 replicates are shown. Red numbers indicate ancestral HPRTs (Anc1 – 7). The tree reveals two main lineages: one with dimer-dimer interaction motifs and one lacking the interaction motifs. See Materials and Methods for tree construction. **B)** Alignment of HPRT dimer-dimer interface from *S. cerevisiae* HPRT, the ancestors (Anc1 – 7) preceding the (p)ppGpp-regulated lineage, and example extant HPRTs. A dimer-dimer interaction motif (shaded gray) is established by Anc7, and all extant HPRTs share Lys81. For the ancestors, red indicates high (>90%), yellow moderate (50-90%), and no color low (<50%) confidence level in the residue identity. Figure Supplements 1 and 2 show the ancestry and residues conserved in the dimeric lineage. **C)** *B. subtilis* K81E and K81A HPRTs have weakened tetramerization according to size-exclusion chromatography. Figure Supplement 3 shows Lys81 at the dimer-dimer interface. **D)** Binding of ^32^P-labeled pppGpp to wild type (black circles), K81A (red squares), and K81E (blue triangles) HPRTs using DRaCALA. Error bars represent SEM of three replicates.

A tracing of the dimer-dimer interface pinpointed residues that coevolve with (p)ppGpp regulation. We followed the change in interface residues from the dimeric Anc1 through Anc7, the ancestor common to the (p)ppGpp-inhibited HPRTs (Figure 5A). We identified a cluster of seven interface residues that changed identity between Anc1 and Anc3 and all seven residues were established by Anc7 (Figure 5B). These residues comprise a β strand and loop that interact with one another across the dimer-dimer interface, so their evolution is likely important for HPRT tetramerization (Figure 5 – figure supplement 3). One residue from this motif, Lys81, had evolved by Anc3 and is conserved in (p)ppGpp-regulated HPRTs but not in (p)ppGpp-insensitive HPRTs (Figure 5B and Figure 5 – figure supplement 1), suggesting that (p)ppGpp regulation is associated with the evolution of this residue. Lys81 reaches across the dimer-dimer interface from each subunit (Figure 5 – figure supplement 3). Therefore, we constructed Lys81 variants with weakened dimer-dimer interfaces (Figure 5C). One variant with a charge reversal (K81E) resulted in a mostly dimeric HPRT and a less disruptive K81A HPRT exhibited a rapid equilibrium between tetramer and dimer (Figure 5C). Importantly, pppGpp bound less well to both K81A and K81E HPRT variants relative to wild type HPRT (Figure 5D). We conclude that the evolution of the HPRT dimer-dimer interface has allowed HPRTs to be regulated by (p) ppGpp by sequestering loop II at the dimer-dimer interface, opening the active site for (p)ppGpp binding (Figure 6).

**Figure 6.**
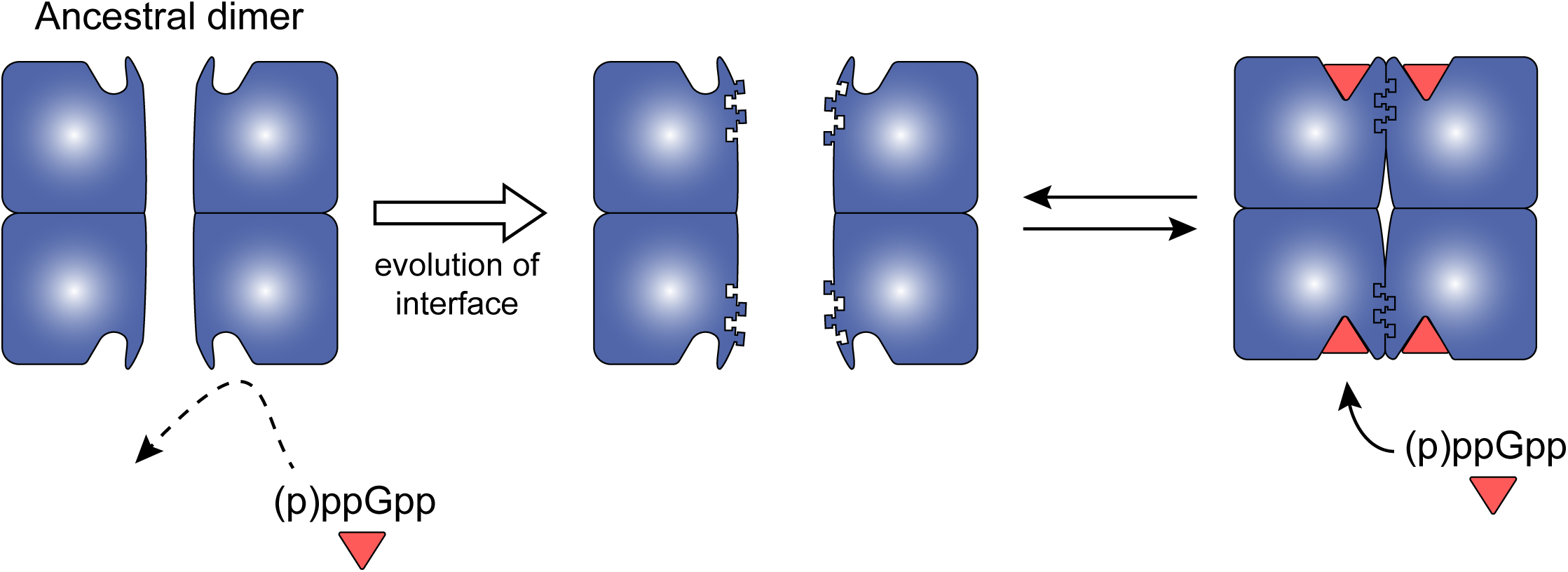
Model for coevolution of protein oligomerization and ligand binding. Bacterial HPRTs have appeared to evolve from a dimeric common ancestor (Figure 5). Dimeric HPRTs interact weakly with (p)ppGpp since the conformation of loop II occludes (p)ppGpp binding (Figure 4). The majority of HPRTs have evolved a dimer-dimer interface motif that sequesters loop II at the interface making the active site conducive to (p)ppGpp binding and inhibition. See Figure Supplement 1 for another possible example of protein-protein interaction affecting ligand binding in the guanylate kinase-(p)ppGpp interaction.

## DISCUSSION

(p)ppGpp is a stress-induced signaling molecule in bacteria that is also critical for cellular fitness and homeostasis even at basal levels. However, (p)ppGpp targets and their evolution in different bacteria remain poorly understood. Here we have described a mechanism explaining how basal levels of (p)ppGpp potently regulate the activity of the housekeeping enzyme HPRT through a novel binding site that may represent a new class of (p)ppGpp effectors. Intriguingly, this site overlaps completely with the active site and is conserved among bacteria, yet differential regulation by (p)ppGpp can be achieved through variation of an allosteric component: the interaction between dimeric subunits of the HPRT tetramer. This interaction tethers a flexible loop at the interface and away from the active site, allowing the binding pocket to accomodate (p)ppGpp. Lack of the dimer-dimer interaction causes HPRT to occlude (p)ppGpp and favor substrate binding. This dimer-dimer interaction is due to an interface motif that appears to have co-evolved with (p)ppGpp binding in the majority of bacterial HPRTs sensitive to (p)ppGpp, whereas dimeric HPRTs without the motif are resistant to (p)ppGpp. We conclude that evolution of the dimer-dimer interface in tetrameric HPRTs has potentiated (p)ppGpp binding to enable basal (p)ppGpp modulation of metabolism, thus increasing bacterial fitness in fluctuating environments. The mechanism of HPRT regulation through tetramerization presents an example of a novel molecular principle of “oligomeric allostery” where oligomerization determines conformations most favorable to ligand binding.

### A novel, high affinity (p)ppGpp binding motif

Our HPRT-ppGpp structure revealed a binding motif distinct from known (p)ppGpp-protein interactions. There are over thirty known (p)ppGpp targets across bacteria, but identifying motifs associated with (p)ppGpp binding has been difficult (Corrigan et al., 2016; Wang et al., 2018; Zhang et al., 2018). In a few cases (p)ppGpp binds allosteric sites at protein interfaces (Kanjee et al., 2011; Ross et al., 2013, 2016; Steinchen et al., 2015; Wang et al., 2018). For many other targets, (p)ppGpp binds at a GTP binding site, leading to overlapping (p)ppGpp and GTP binding motifs (Fan et al., 2015; Kihira et al., 2012; Liu et al., 2015b; Pausch et al., 2018; Rymer et al., 2012). In the case of HPRT, however, the (p)ppGpp binding site is not at an interface nor is it a GTP binding site. Instead, it shares a well-characterized motif associated with PRPP binding (EDIIDSGLT in *B. anthracis* Hpt-1) (Sinha and Smith, 2001). Other PRPP-binding proteins, including UPRT and APRT, have been shown to bind (p)ppGpp (Wang et al., 2018; Zhang et al., 2018), and it is likely that (p)ppGpp also binds to their PRPP motif. Identification of this motif may provide a new class of (p)ppGpp-binding proteins, allowing us to predict additional targets.

(p)ppGpp interacts with proteins in two main conformations: elongated, with the phosphate arms extended away from one another in a T shape, and ring-like, with the phosphate arms near one another in a Y shape. The compact, ring-like conformation has been associated with higher-affinity interactions than the elongated conformation (Steinchen and Bange, 2016). However, in the HPRT-ppGpp interaction, ppGpp takes an elongated conformation, but exhibits strikingly tight affinities as high as K_d_ ≈ 0.1 μM for some species (Figure 1F and Table 1). This high affinity may be due to extensive backbone amide interactions with both phosphate arms as well as (p)ppGpp’s close mimicry of substrate binding (Figure 2D). It is likely that higher affinity interactions with the elongated (p)ppGpp conformation will become more common as additional targets are characterized.

### HPRT tetramerization enables basal (p)ppGpp inhibition

Our data show that the dimer-dimer interface of HPRT potentiates (p)ppGpp regulation by allosterically promoting a conformation conducive to ligand binding (Figure 6). HPRT’s tetramerization harnesses the flexible loop II to open the binding pocket (Figure 4A) for (p)ppGpp to bind with high affinity. Loop II’s influence on (p)ppGpp binding may be the reason that it has evolved to be part of the interface of bacterial HPRTs, whereas in eukaryotic (e.g. human) HPRTs, which likely do not interact with (p)ppGpp in nature, loop II is on the outside of the oligomer facing solvent (Eng et al., 2015).

Our model can also explain the evolutionary significance of bacterial HPRTs functioning as dimers without dissociating to monomers. Strength of protein-protein interactions has been correlated with increased buried surface area at the interface (Nooren and Thornton, 2003). In a hypothetical monomer-monomer interaction, the smaller loop II interface may be too weak and too transient to keep loop II away from the active site. In the interaction between two homodimers, the loop II interface is duplicated, which increases the surface area and provides anchor points to hold loop II at the interface and away from the (p)ppGpp binding pocket (Figure 6).

Our characterization of HPRT explains how bacteria can be regulated by basal levels of (p)ppGpp. While (p)ppGpp is mostly chacterized as a regulator of gene expression that functions in starvation-induced concentrations, basal concentrations of (p)ppGpp are known to be responsible for sustaining antibiotic tolerance and virulence in *E. faecalis* (Gaca et al., 2013), maintaining cyanobacterial light/dark cycles (Puszynska and O’Shea, 2017), influencing rRNA expression and growth rate in *E. coli* (Potrykus et al., 2011), and regulating GTP synthesis in Firmicutes (Gaca et al., 2013; Kriel et al., 2012). However, the principles that determine how basal and induced (p)ppGpp regulate different cellular targets remained unclear. In the single species *B. subtilis*, for example, (p)ppGpp interacts with DNA primase (K_i, ppGpp_ = 250 μM), IMP dehydrogenase (K_i, ppGpp_ = 50 μM), and guanylate kinase (K_i, pppGpp_ = 14 μM), allowing them to be inhibited *in vivo* at induced (p)ppGpp levels (Liu et al., 2015b; Pao and Dyess, 1981; Wang et al., 2007). (p)ppGpp’s interaction with *B. subtilis* HPRT is far stronger (K_d_ = 0.75 μM; Table 1), potentially allowing (p)ppGpp to modulate its activity at basal levels throughout growth. For some species, such as *Bacteroides* spp., where HPRT is the only known target of (p)ppGpp, the K_d (pppGpp)_ for HPRT is as low as 0.1 μM (Figure 1F and Table 1). Our data suggest that the dimer-dimer interaction is a mechanism that greatly strengthens (p)ppGpp binding affinity, allowing basal (p)ppGpp to regulate HPRT and thus constantly regulate purine nucleotide synthesis via the salvage pathway.

The evolution of HPRT inhibition may be driven in part by environmental niche. For example, bacteria from the mammalian intestinal tract have evolved the dimer-dimer interface motif (Figure 5) and have highly (p)ppGpp-sensitive HPRTs (Figure 1), possibly to robustly maintain intracellular metabolism despite fluctuations in exogenous purines that could depend on the purine content of the diet (Choi et al., 2004; Kaneko et al., 2014; Zgaga et al., 2012). On the other hand, HPRTs that have not evolved the dimer-dimer interface motif share an intracellular niche, including obligate (*C. burnetii*) and facultative (*L. pneumophila*, *P. aeruginosa*, *N. meningitidis*) intracellular pathogens that salvage purines inside their host cell (Ducati et al., 2011; Miller and Thompson, 2002; Traut, 1994a). Reduced regulation by (p)ppGpp may enable salvage and nucleotide synthesis in conditions where (p)ppGpp levels are elevated.

### Protein oligomerization allosterically alters ligand specificity

The mechanism we characterized for HPRT may represent a more broadly applicable principle by which protein oligomerization changes the conformation of a ligand binding pocket to alter ligand specificity. Oligomerization of proteins into homomers can provide multiple adaptive advantages, including mechanisms of allosteric regulation, promoting protein stability, providing complete active sites or ligand binding sites at oligomeric interfaces, maintaining proximity of signal transduction, and serving cytoskeletal or other structural roles (Ali and Imperiali, 2005; Matthews and Sunde, 2012; Perica et al., 2012; Traut, 1994b). Oligomerization of metabolic enzymes, for example, is prevalent for allosteric enzyme regulation, where ligand binding to a regulatory site causes oligomeric alterations to facilitate conformational changes at an active site (Traut, 2008). In the case of HPRT, however, oligomerization into higher order tetramers is not shaped by the requirement of a canonical allosteric regulation because (p)ppGpp does not bind to a different site than the active site. Nor does the oligomeric state of HPRT affect the protein stability or activity, which are common oligomeric adaptive advantages (Figure 4 – figure supplement 3 and Table 4). Instead, HPRT tetramerization allows strong inhibition by basal (p)ppGpp, suggesting that the adaptive advantage of tetramerization is altering (p)ppGpp specificity for enhanced regulation by this nucleotide. There are numerous examples of ligands stabilizing protein oligomerization (Traut, 1994b). But we have identified an example of protein oligomeric state dictating the binding of a ligand at a non-interface binding site by allosterically inducing changes in the ligand binding pocket (Figure 4F), thus constituting a new principle of “oligomeric allostery.”

**Table 4.**
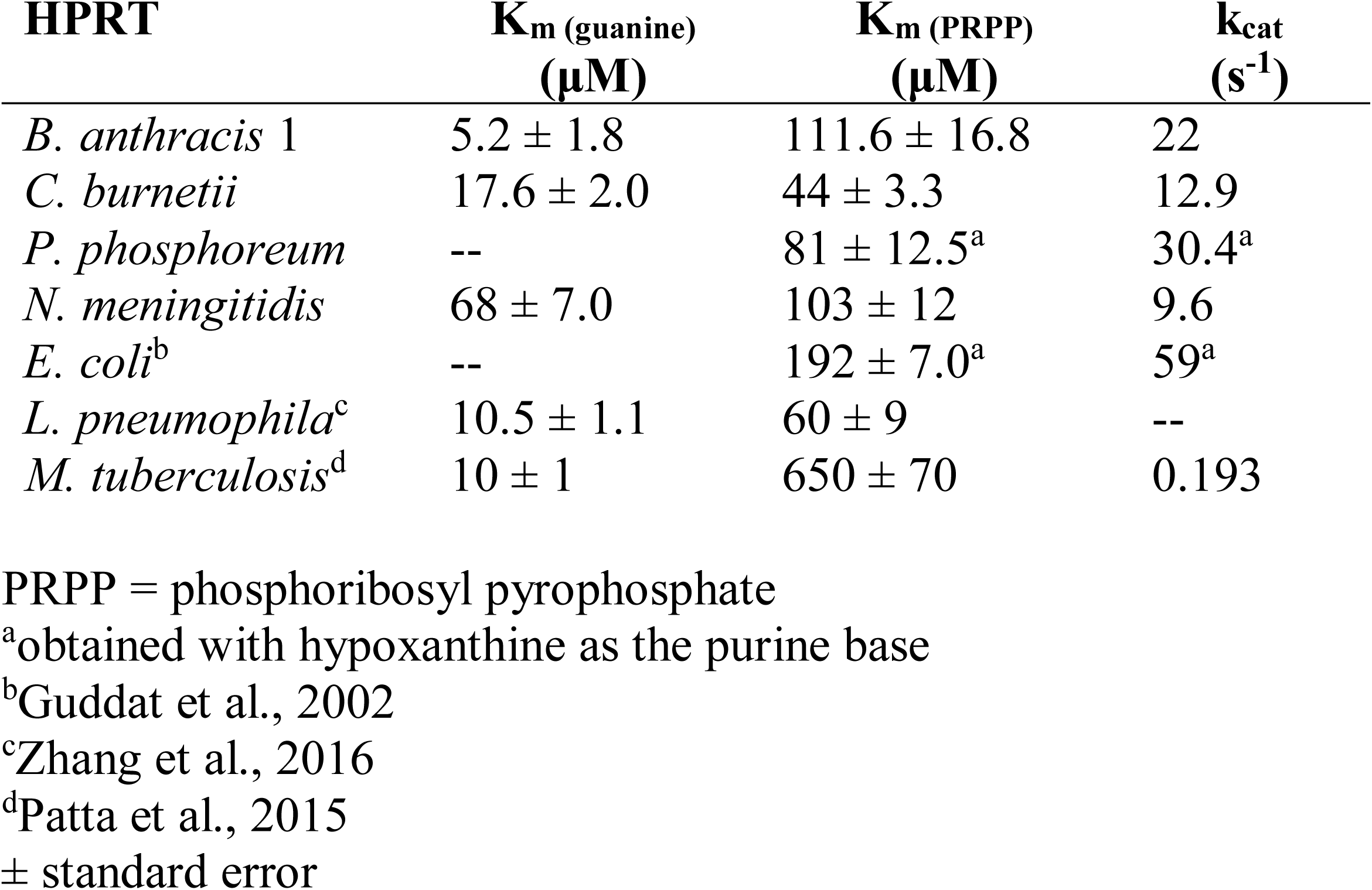
HPRT kinetic parameters.

| HPRT | $K_m$ (guanine)<br>($\mu\text{M}$) | $K_m$ (PRPP)<br>($\mu\text{M}$) | $k_{cat}$<br>($\text{s}^{-1}$) |
| --- | --- | --- | --- |
| *B. anthracis* 1 | $5.2 \pm 1.8$ | $111.6 \pm 16.8$ | 22 |
| *C. burnetii* | $17.6 \pm 2.0$ | $44 \pm 3.3$ | 12.9 |
| *P. phosphoreum* | -- | $81 \pm 12.5^a$ | $30.4^a$ |
| *N. meningitidis* | $68 \pm 7.0$ | $103 \pm 12$ | 9.6 |
| *E. coli*<sup>b</sup> | -- | $192 \pm 7.0^a$ | $59^a$ |
| *L. pneumophila*<sup>c</sup> | $10.5 \pm 1.1$ | $60 \pm 9$ | -- |
| *M. tuberculosis*<sup>d</sup> | $10 \pm 1$ | $650 \pm 70$ | 0.193 |

PRPP = phosphoribosyl pyrophosphate
<sup>a</sup>obtained with hypoxanthine as the purine base
<sup>b</sup>Guddat et al., 2002
<sup>c</sup>Zhang et al., 2016
<sup>d</sup>Patta et al., 2015
± standard error

Oligomeric allostery may play a role in altering specificity for other (p)ppGpp protein targets across bacteria. For example, we previously found that the enzyme guanylate kinase (GMK) is inhibited by (p)ppGpp in multiple phyla of bacteria but not in Proteobacteria despite the fact (p)ppGpp binds the conserved active site (Liu et al., 2015b). This puzzle may now be explained by oligomeric allostery: GMK is a dimer. In the (p)ppGpp-sensitive GMK, a lid domain opens the active site for (p)ppGpp binding through pulling of the lid by the C-terminal helix of the adjoining monomer in a GMK dimer (Figure 6 – figure supplement 1). In the (p)ppGpp-insensitive *E. coli* GMK, the lid domain is closed, with the C-terminal helix from the adjoining monomer perhaps responsible for stabilizing the closed position (Figure 6 – figure supplement 1). It remains to be tested whether the monomer-monomer interaction in GMK influences the lid domain conformation and (p)ppGpp binding.

It is striking to note that a majority of ligands bind within 6 Å of a protein-protein interface (Gao and Skolnick, 2012). Thus this principle of “oligomeric allostery” likely extends to many other protein-ligand interactions.

### Coevolution of ligand binding and protein oligomerization

One potential advantage of oligomeric allostery is that it provides evolutionary flexibility for ligand binding. Evolving different residues within ligand binding sites can alter ligand specificity, but active sites that bind substrates and inhibitors are under functional and evolutionary constraints to maintain enzymatic activity (Echave et al., 2016; Huang et al., 2015). On the other hand, protein-protein interfaces are more evolutionarily flexible, particularly when they are not obligate interactions like the dimer-dimer interface of HPRT (Echave et al., 2016; Mintseris and Weng, 2005). This suggests that changing oligomeric states could be an evolutionarily flexible mechanism for altering ligand specificity (Figure 6). Indeed, the coevolution of (p)ppGpp regulation and the HPRT dimer-dimer interaction has provided us with an example of how protein oligomerization and ligand binding can coevolve, demonstrating that organisms have already adopted this strategy.

Many proteins are regulated by small molecules (Gao and Skolnick, 2012; Najmanovich, 2017). Our results suggest that ligand binding, even at non-interface binding pockets, influence evolutionary diversification of protein oligomers potentially through purifying selection of conformations that favor protein-ligand interactions. While homomeric evolution of some proteins has been implicated as physicochemical or stochastic processes (Abrusán and Marsh, 2018; André et al., 2008; Lukatsky et al., 2007; Lynch, 2013), our data provide evidence for ligand binding as an adaptive advantage driving the evolutionary diversification of protein homomers. Given the proximity of ligand binding sites to protein interfaces (Gao and Skolnick, 2012), and since it is easier to evolve protein-protein interactions (Perica et al., 2012) rather than evolving new sites for allosteric regulation, such an adaptive benefit is likely to exist more broadly beyond HPRT and (p)ppGpp in other protein-ligand interactions.

## MATERIALS AND METHODS

### Plasmid construction and mutagenesis

For purification and DRaCALA analyses, *hprT* coding sequences were cloned into the pLIC-trPC-HA vector (pJW269) using the ligation independent cloning (LIC) protocol (Eschenfeldt et al., 2009; Stols et al., 2002) and was scaled up to include more *hprT* sequences when necessary as described (Abdullah et al., 2009). Bacterial *hprT* homologs were amplified from genomic DNA, cloned using LIC, and transformed into *E. coli* BL21(DE3) (see Supplementary File 1 for primers and plasmids). Amino acid substitutions were performed either with QuikChange XL Site-Directed Mutagenesis (Agilent Technologies) or with a megaprimer site-directed mutagenesis protocol (Kirsch and Joly, 1998). Loop II deletions were made using a protocol as described (Hansson et al., 2008). pLIC-trPC-HA inserts were amplified and sequenced for confirmation using oJW1124 and oJW492.

### Protein purification

HPRTs were recombinantly expressed in *E. coli* BL21(DE3) (NEB) from a pLIC-trPC-HA plasmid with the gene inserted downstream of a sequence encoding a 6X histidine tag and a tobacco etch virus (TEV) protease recognition site. Seed cultures grown to mid-log phase were diluted 1:50 into batch culture of LB supplemented with 100 μg/mL carbenicillin. Protein synthesis was induced at OD_600_ ≈ 0.8 with 1 mM IPTG for four hours. Cells were pelleted and stored at −80° C until purification.

For large scale purifications, cells were resuspended in Lysis Buffer (50 mM Tris-HCl pH 7.5, 500 mM NaCl, 10 mM imidazole) and lysed with a French press. The lysate was centrifuged to obtain the soluble fraction, which was filtered through 0.45 μm filters. The sample was put over a HisTrap FF column (GE Healthcare) on an AktaPure FPLC (GE Healthcare), the column was washed with 20 column volumes of Lysis Buffer, and the protein was eluted with a gradient of increasing Elution Buffer (50 mM Tris-HCl pH 7.5, 500 mM NaCl, 250 mM imidazole). Recombinant protein fractions were dialyzed with 10 mM Tris-HCl (pH 7.5), 100 mM NaCl, 1 mM DTT, and 10% glycerol prior to concentrating and flash-freezing for storage. For small scale purifications, Ni-NTA spin columns were used according to manufacturer’s instructions (Qiagen). Lysis buffer was 100 mM sodium phosphate (pH 8.0), 500 mM NaCl, and 10 mM imidazole. Wash and elution buffers were the same as the lysis buffer except with 20 mM and 500 mM imidazole, respectively. Protein purity was determined using SDS-PAGE, and concentrations were measured using the Bradford assay (Bio-Rad) or using A_280_ with extinction coefficients calculated by ProtParam (SIB ExPASy Bioinformatics Resource Portal).

To purify 35 HPRT homologs for activity assays (see Figure 1), a 96-well Capturem His-tagged purification kit (Clontech) was used according to the manufacturer’s instructions. The following buffers were used: lysis [xTractor (Clontech) + 1 μg/mL DNase I], wash [20 mM Na_3_PO_4_ pH 7.6, 200 mM NaCl, 10 mM imidazole], elution [20 mM Na_3_PO_4_ pH 7.6, 500 mM NaCl, 500 mM imidazole]. Following purification, the buffer was exchanged to 10 mM HEPES pH 8, 150 mM NaCl, 10 mM MgCl_2_, 1 mM DTT using Zeba spin desalting plates (Thermo Scientific) according to the manufacturer’s instructions. The Bradford assay was used to measure protein concentration, and the proteins were aliquoted at 8 μM and flash-frozen with liquid nitrogen.

For crystallography with ppGpp, recombinant *B. anthracis* Hpt-1 was purified as described above followed by dialysis with His-tagged TEV protease in 10 mM Tris-HCl (pH 7.5), 100 mM NaCl, and 1 mM DTT. The dialyzed protein was incubated with Ni-NTA beads for 30 minutes, the beads were centrifuged and the supernatant was run over a HiPrep Sephacryl 16/60 S-100 HR column (GE Healthcare) in Crystal Buffer 1 (10 mM Tris-HCl pH 8.3, 250 mM NaCl, and 1 mM DTT). Relevant fractions were further dialyzed in the Crystal Buffer 1 prior to concentrating to ≈10 mg/mL. For crystallography with substrates, Crystal Buffer 2 was used (10 mM Tris-HCl pH 8 and 100 mM NaCl).

### Enzyme inhibition assays

HPRT activity assays were performed as described previously (Biazus et al., 2009; Kriel et al., 2012; Xu et al., 1997). The standard assay was performed at 25° C and contained 100 mM Tris-HCl (pH 7.4), 12 mM MgCl_2_, 1 mM PRPP (MilliporeSigma), 50 μM guanine or hypoxanthine (MilliporeSigma), and 20 nM HPRT. Reactions were initiated with the purine base and monitored in a spectrophotometer (Shimadzu UV-2401PC) at 257.5 nm or 245 nm for conversion of guanine to GMP or hypoxanthine to IMP, respectively. A difference in extinction coefficients of 5900 M^-1^cm^-1^ was used for GMP and guanine and 1900 M^-1^cm^-1^ for IMP and hypoxanthine. For inhibition curves, assays were performed at the substrate concentrations listed above and at variable pppGpp, ppGpp, and pGpp concentrations. (p)ppGpp was synthesized as described (Liu et al., 2015b). Initial velocities of the inhibited reactions were normalized to the uninhibited initial velocity prior to fitting to the equation Y = 1/(1 + (x / IC_50_)^s^) to calculate IC_50_.

To test inhibition of HPRT homologs (see Figure 2A), reactions were performed in a Synergy 2 microplate reader (BioTek). The assay was performed at 25° C with 50 μM hypoxanthine, 1 mM PRPP, and 100 nM HPRT in the same reaction buffer as above. For *Coxiella burnetii* HPRT, 50 μM guanine was used as the substrate since its activity was very low with hypoxanthine. Reactions were performed in triplicate without (p)ppGpp, with 25 μM ppGpp, and with 25 μM pppGpp. Reaction rates from the first-order kinetic curves were determined using R (v 3.4.3).

### Isothermal titration calorimetry

Experiments were performed using the MicroCal iTC_200_ (GE Healthcare). *B. anthracis* Hpt-1 was dialyzed into the ITC buffer (10 mM HEPES pH 8, 150 mM NaCl, 10 mM MgCl_2_) with three buffer changes at 4° C. The concentration of protein was calculated using a molar extinction coefficient of 16390 M^-1^cm^-1^ and A_280(true)_ (A_280_ – (1.96 × A_330_) (Pace et al., 1995). The experiments were performed at 25° C with 45.5 μM HPRT, a reference power of 6 μCal/s, and a stirring speed of 1000 RPM. pppGpp was solubilized in dialysate from protein dialysis and its concentration was measured using a molar extinction coefficient of 13700 M^-1^cm^-1^ and A_253_. pppGpp was titrated into *B. anthracis* Hpt-1 with the following: 1×1 μL (discarded), 19×2 μL. Data analysis and one-site binding modelling (where relevant) was performed using MicroCal Origin 5.0 software provided by the company.

### X-ray crystallography

Proteins were prepared for crystallography as described above. For crystals formed with pppGpp, *B. anthracis* Hpt-1 was concentrated to ~10 mg/mL in 10 mM Tris pH 8.3, 250 mM NaCl, and 1 mM DTT. The pppGpp ligand was resuspended in ddH_2_O. MgCl_2_ was added to the protein at a final concentration of 1 mM and crystals were formed using hanging drop vapor diffusion with 900 μL of reservoir liquid in each well. Each drop contained 0.9 μL protein, 0.9 μL reservoir liquid, and 0.2 μL pppGpp (final concentration, 1.5 mM pppGpp). Crystals formed in 0.2 M ammonium tartrate dibasic pH 6.6 and 20% PEG 3350 in 3-6 months. Identical crystals formed in replicated conditions in 1-2 weeks. Crystals were soaked in reservoir liquid with 25% glycerol prior flash freezing in liquid nitrogen. Apo protein crystals formed within 1-2 days in multiple conditions with high sulfate concentrations. Using protein preparations from above, apo crystals formed in 0.01 M CoCl_2_, 0.1 M MES monohydrate pH 6.5, 1.8 M ammonium sulfate, and crystals were soaked in reservoir liquid with 25% glycerol for cryoprotection prior to freezing.

For crystals with substrates, *B. anthracis* Hpt-1 was concentrated to ~10 mg/mL in 10 mM Tris pH 8 and 100 mM NaCl. PRPP was resuspended in ddH_2_O and 9-deazaguanine was resuspended in 100% DMSO. Additives were diluted in the protein solution at final concentrations of 10 mM MgCl_2_, 2 mM PRPP, and 1 mM 9-deazaguanine prior to crystallization. Drops for hanging drop vapor diffusion comprised 1 μL crystal condition and 1 μL protein/ligand mixture. Crystals formed within 3 days in 0.2 M ammonium acetate, 0.1 M sodium acetate trihydrate pH 4.6, 30% PEG 4000. Reservoir solution with 25% ethylene glycol was added to the drops for cryoprotection prior to flash freezing in liquid nitrogen.

Diffraction data was collected at the Life-Science Collaborative Access Team (LS-CAT), beamline 21-ID-F (Hpt-1 with sulfates), 21-ID-G (Hpt-1 with ppGpp), and 21-ID-D (Hpt-1 with substrates) at the Advanced Photon Source (APS) at Argonne National Labs (Argonne, IL). Data was indexed and scaled using HKL2000 (Otwinowski and Minor, 1997). Phasing for Hpt-1 with sulfates and ppGpp was determined by molecular replacement with PDB ID 3H83 as a search model using Phenix (Adams et al., 2010). Phasing for Hpt-1 with substrates was determined by molecular replacement with Hpt-1-ppGpp as a search model using Phenix. Iterative model building with Coot and refinement with Phenix produced final models (Emsley and Cowtan, 2004).

### Size exclusion chromatography

Size exclusion chromatography was performed using AktaPure and a Superose 12 10/300 GL column (GE Healthcare, Inc.). A buffer consisting of 10 mM HEPES pH 8.0, 100 mM NaCl, and 10 mM MgCl_2_ was used to run ≈10 – 20 μM (0.2 – 0.5 mg/mL) protein over the column at a rate of 0.1 – 0.25 mL/min. Additional 1 mM DTT was used for proteins containing cysteines. For gel filtration with ligands, 500 μM PRPP or 500 μM pppGpp was included in the buffer. A gel filtration standard (Bio-Rad) was used to establish molecular weight, and bovine serum albumin (BSA) was included as an additional marker.

### Dimethyl adipimidate crosslinking

Crosslinking was performed with ≈10 μM *B. subtilis* HPRT, 20 mM dimethyl adipimidate (DMA) (ThermoFisher), and 500 μM ligand. DMA was suspended in 25 mM HEPES pH 8.0, 100 mM NaCl, 10 mM MgCl_2_, and 10% glycerol, and the solution was buffered to pH 8.5. HPRT was dialyzed into 25 mM HEPES pH 7.5, 100 mM NaCl, and 10% glycerol. Ligands were incubated with protein for 10 minutes followed by a 15 minute incubation with DMA at room temperature. Reactions were terminated with addition of 2X Laemmli buffer (Bio-Rad) for immediate analysis with SDS-PAGE (10% polyacrylamide gel). Gels were stained with SYPRO Ruby (Bio-Rad) according to manufacturer’s protocol and imaged using a Typhoon FLA9000 (GE Healthcare).

### Dynamic light scattering

Dynamic light scattering was performed using DynaPro99 (Protein Solutions/Wyatt Technologies). Readings were from a 20 μL solution (10 mM Tris-HCl pH 8, 100 mM NaCl, 10 mM MgCl_2_, 6.5% glycerol) with 4 mg/mL (≈175 μM) *B. anthracis* Hpt-1 and 2 mM ligands. Data were collected and analyzed using Dynamics version 5.25.44 software.

### DRaCALA

DRaCALA was performed with pure protein and radioactive ligand as described (Roelofs et al., 2011). [3′ -α ^32^P] pppGpp was synthesized according to modified protocols of non-radioactive and radioactive pppGpp syntheses (Corrigan et al., 2016; Hogg et al., 2004; Mechold et al., 2002). The reaction contained 25 mM bis-Tris propane (pH 9.0), 15 mM MgCl_2_, 0.5 mM DTT, 2 mM ATP, 2 μM Rel_Seq_, and 37.5 μCi [α-^32^P] GTP (Perkin Elmer). The reaction was incubated at 37 °C for one hour. The reaction was diluted in 0.5 mL of Buffer A (0.1 mM LiCl, mM EDTA, 25 mM Tris-HCl pH 7.5) prior to adding to a HiTrap QFF (1 mL) strong anion exchange column (GE Healthcare) equilibrated with 10 column volumes (CV) of Buffer A. The column was washed with 10 CV of Buffer A followed by an additional wash with 10 CV of 83% Buffer A + 17% Buffer B (Buffer B: 1 M LiCl, 0.5 mM EDTA, 25 mM Tris-HCl pH 7.5). ^32^P-pppGpp was eluted with a mixture of 50% Buffer A + 50% Buffer B. Fractions of 1 mL were collected from the elution. For DRaCALA, a final dilution of 1:200 of the first fraction was typically used.

DRaCALA reactions (20 μL) using purified HPRT contained 100 mM Tris-HCl (pH 7.4), 12 mM MgCl_2_, 10 μM protein, and ^32^P-pppGpp. Protein was dialyzed or diluted into buffer lacking glycerol, as glycerol interferes with diffusion of the aqueous phase. Reactions were incubated at room temperature for 10 minutes. Two microliters from each reaction were spotted in duplicate on Protran BA85 nitrocellulose (GE Healthcare). Spots were allowed to dry and radioactivity was detected with phosphorimaging (Typhoon FLA9000). Fraction bound of ^32^P-pppGpp was calculated as described (Roelofs et al., 2011). Data were analyzed in GraphPad Prism v5.02 and fitted to the equation Y = (B_max_ × K_d_) / (K_d_ + X) (Roelofs et al., 2011).

DRaCALA using cell lysates was adapted from a previous protocol (Roelofs et al., 2015). One milliliter of cells containing overexpressed recombinant HPRT was pelleted and resuspended in 100 μL of binding buffer (20 mM Tris pH 8, 100 mM NaCl, 12 mM MgCl_2_, 0.1 mM DTT) supplemented with 1 μM PMSF, 250 μg/mL lysozyme, and 10 μg/mL DNase I. Cells were lysed with freeze/thaw cycles. In a 20 μL DRaCALA reaction, 10 μL of cell lysate was added to binding buffer and ^32^P-pppGpp. Reactions were performed and analyzed as above. For measuring binding affinity (K_d_) of proteins in cell lysates, recombinant protein expression level in cell lysates was determined by comparing expression to a standard of purified *B. subtilis* HPRT co-resolved with SDS-PAGE.

### Differential scanning fluorimetry

Differential scanning fluorimetry was performed using 10 μM protein in a buffer containing 20 mM HEPES pH 8, 100 mM NaCl, 10 mM MgCl_2_, 1 mM DTT, and 10X SYPRO orange dye (diluted from 5000X stock; Sigma-Aldrich). Proteins were mixed in an optically clear quantitative PCR (qPCR) 96-well plate and sealed with plastic film. Relative fluorescence intensity was monitored in a Bio-Rad qPCR machine using FRET detection over a temperature increase of 1 °C/min from 25 to 90 °C. Data were analyzed using GraphPad Prism.

### Protein sequence analysis and ancestral sequence reconstruction

Protein sequences were aligned using MUSCLE with default settings in MEGA v7.0. The maximum likelihood phylogenetic tree was constructed using MEGA v7.0 with 1000 bootstrap replications and the Whelan And Golden (WAG) substitution model, assuming gamma distributed substitution rates with invariant sites. Gaps found in fewer than 10% of the sequences were ignored. The HPRT1 sequence from *Saccharomyces cerevisiae* was used as the outgroup. Ancestral HPRT sequences were reconstructed using MEGA X with the phylogenetic tree, maximum likelihood statistics, and the Jones-Taylor-Thornton model (Jones et al., 1992).

## ACKNOWLEDGMENTS

We thank members of the Wang lab for feedback on this manuscript. We thank Federico Rey, Bob Kerby, JD Sauer, Dan Pensinger, Helen Blackwell, and Kayleigh Nyffeler for assistance in obtaining bacterial genomic DNA. We thank Katrina Forest for assistance with dynamic light scattering. The work of J. D. W. was supported in part by a Faculty Scholar grant from the Howard Hughes Medical Institute and by R35 GM127088 from the NIH. B. W. A. was supported by NSF GRFP DGE-1256259.

## AUTHOR CONTRIBUTIONS

J.D.W. conceptualized the study and supervised the research. B.W.A., K.L., and J.D.W. designed the experiments. B.W.A. and K.L. conducted experiments. C.W., K.D., K.A.S., and J.L.K. performed structural analyses. B.W.A., C.W., K.D., J.L.K, and J.D.W. discussed the results and interpretations. B.W.A. prepared the manuscript. B.W.A. and J.D.W. wrote the paper.

## Figure supplements

**Figure 1 - figure supplement 1.**
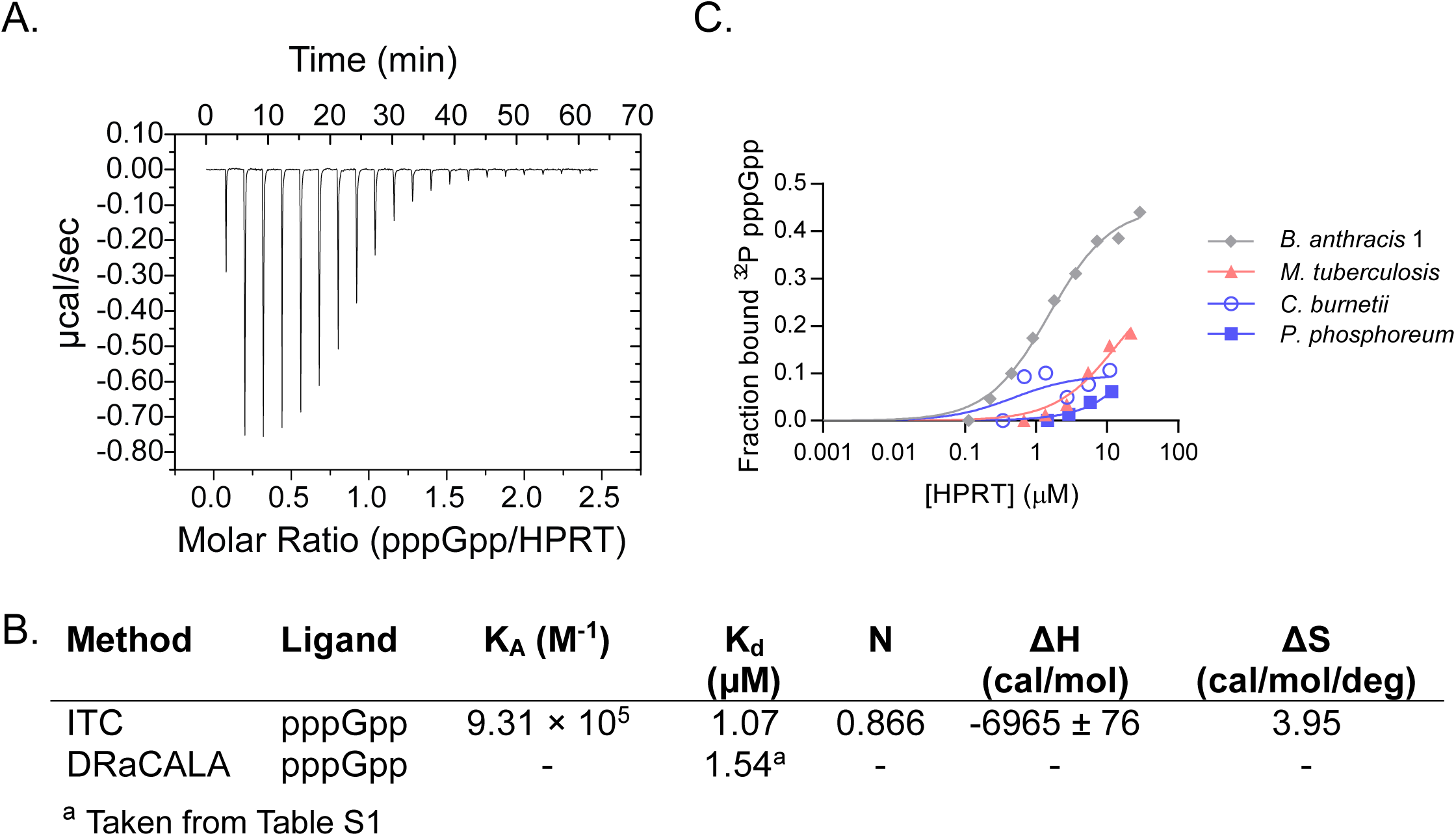
Characterizing pppGpp interaction with bacterial HPRTs. **A)** Energy isotherm of isothermal titration calorimetry between pppGpp and *B. anthracis* Hpt-1. **B)** Parameters from isothermal titration calorimetry and DRaCALA with *B. anthracis* Hpt-1. DRaCALA K_d_ estimated from data in panel C. **C)** DRaCALA binding curves between HPRT and ^32^P-labeled pppGpp performed with a single replicate. The interactions with *M. tuberculosis*, *C. burnetii*, and *P. phosphoreum* HPRTs were too weak to use for a K_d_ approximation. *B. anthracis* Hpt-1 is shown for comparison.

**Figure 2 - figure supplement 1.**
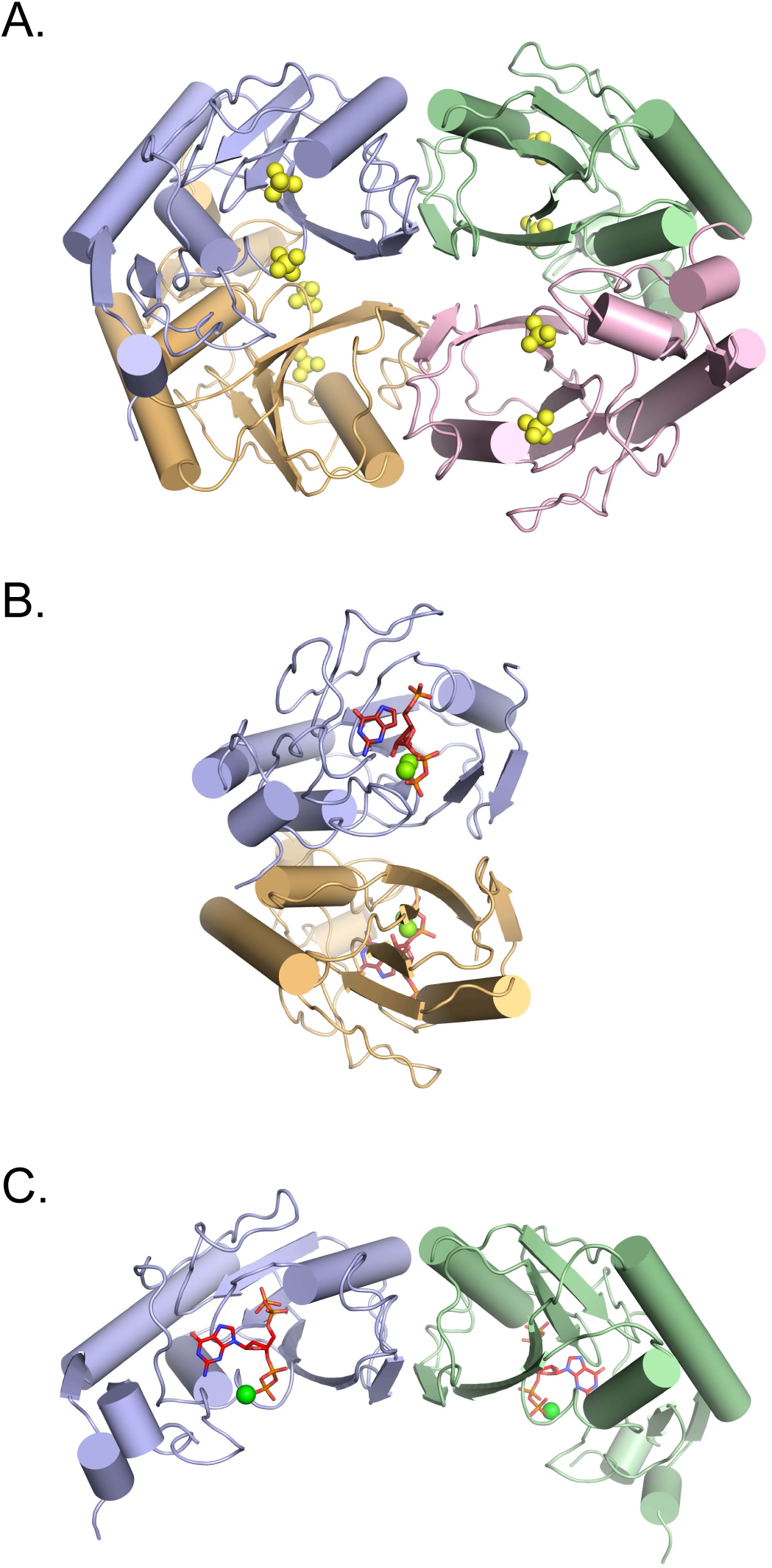
Asymmetric units of *B. anthracis* Hpt-1 structures. **A)** *B. anthracis* Hpt-1 crystallized as a tetramer in the asymmetric unit without ligands. Each color represents a separate monomer. Yellow spheres represent two sulfates bound to each monomer. **B)** *B. anthracis* Hpt-1 crystallized with substrates (PRPP and 9-deazaguanine) as a dimer in the asymmetric unit. Red sticks represent the substrates and the green spheres represent Mg^2+^. **C)** *B. anthracis* Hpt-1 crystallized with ppGpp as a dimer in the asymmetric unit. Red sticks represent pppGpp and the green sphere represents Mg^2+^. Two asymmetric units form a tetramer nearly identical to A.

**Figure 2 - figure supplement 2.**
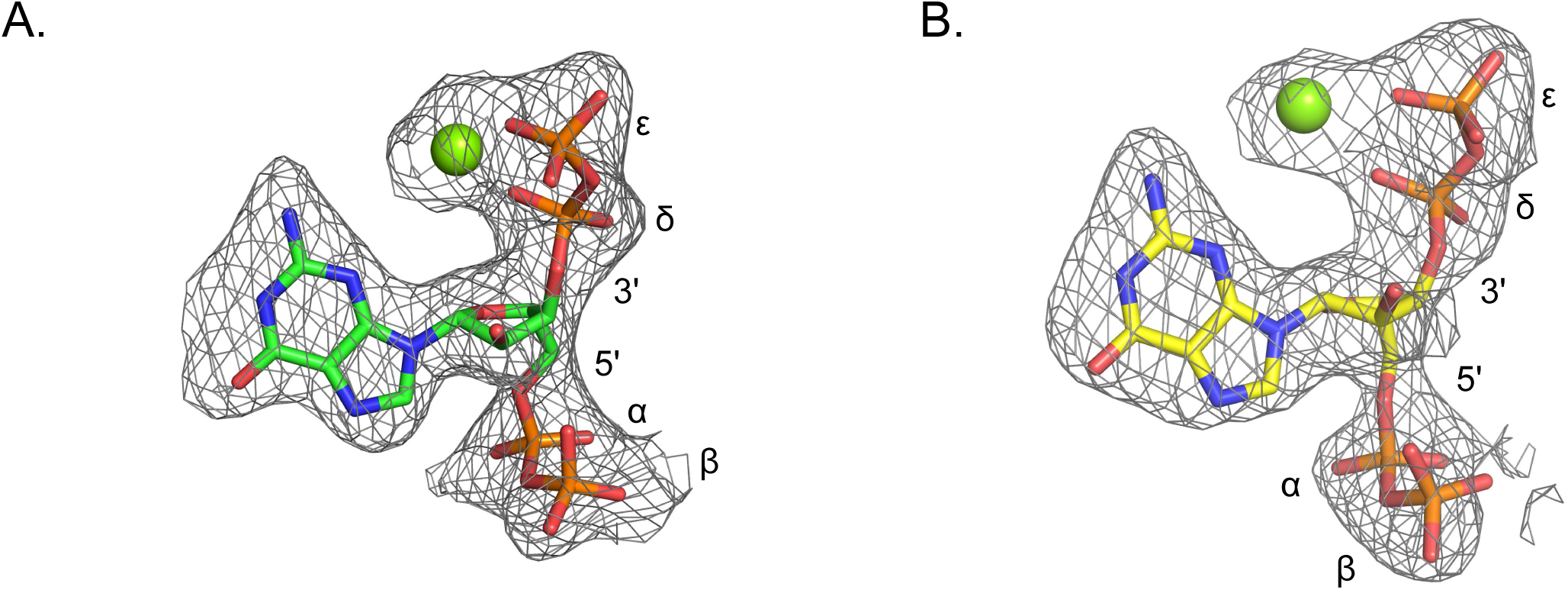
Omit electron densities of ppGpp crystallized with *B. anthracis* Hpt-1. Omit electron densities contoured to 2σ of ppGpp crystallized with *B. anthracis* Hpt-1. ppGpp associated with molecule A of the asymmetric unit is shown in (A) and ppGpp associated with molecule B of the asymmetric unit is shown in (B). Green spheres represent Mg^2+^. Phosphates are labeled α through ε (α and β on 5′, δ and ∈ on 3′).

**Figure 2 - figure supplement 3.**
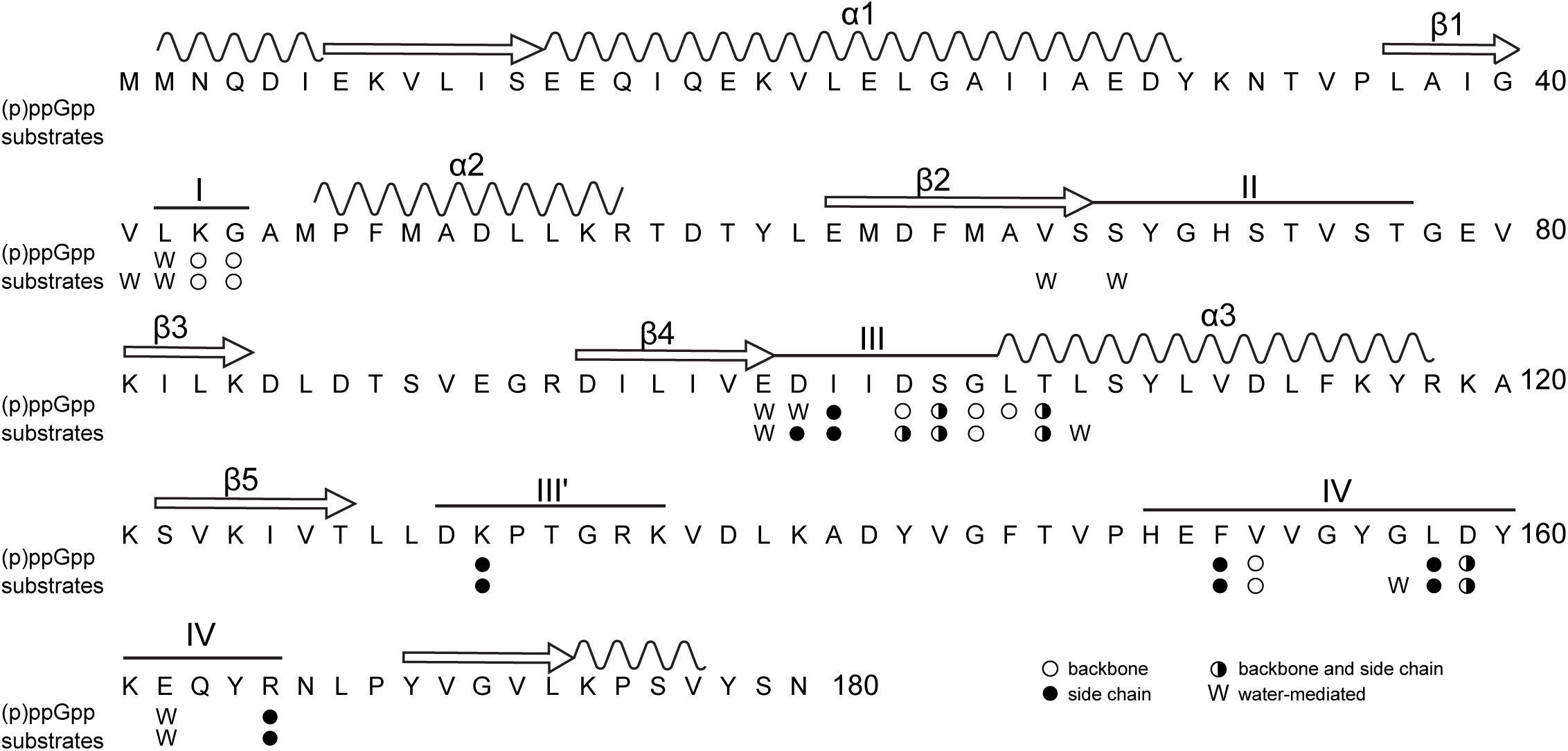
Primary structure of *B. anthracis* Hpt-1 showing ligand-interacting residues. Primary structure of *B. anthracis* Hpt-1 with secondary structure elements annotated above (α1 – 3, β1 – 5, loops I – IV). Residues involved in (p)ppGpp and substrate binding are marked: empty circles denote residues with a backbone interaction with ligands; filled circles denote residues with a side chain interaction; half-filled circles represent residues with both backbone and side chain interactions; and “W” represents water-mediated interactions.

**Figure 2 - figure supplement 4.**
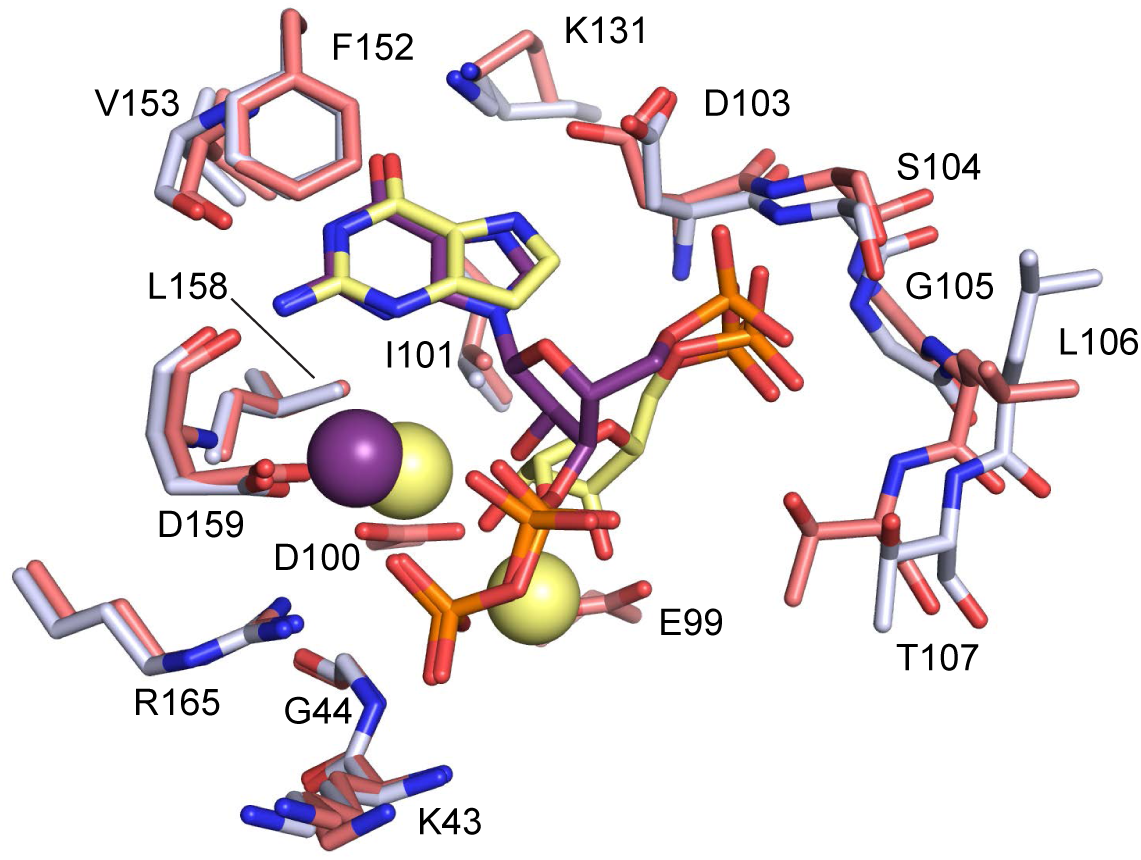
Comparison between ppGpp-bound and substrates-bound binding pocket. Overlay of the active site of ppGpp-bound HPRT (silver) and substrates-bound HPRT (salmon). ppGpp is shown in purple and substrates are shown in yellow. Spheres represent Mg^2+^. E99 and D100, which interact with substrates, are not shown for ppGpp. Val67 and Ser69 on loop II, which interact with PRPP, are omitted.

**Figure 2 - figure supplement 5.**
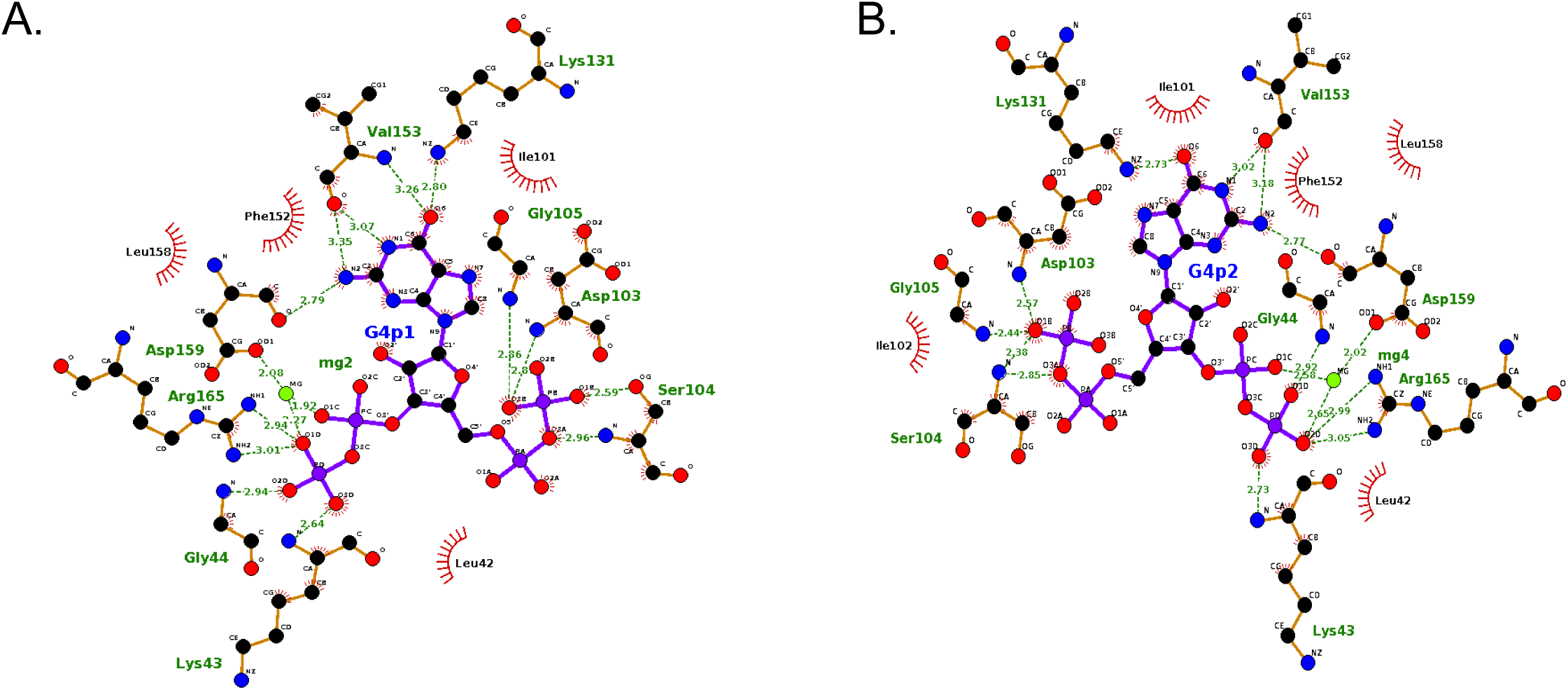
LigPlots of ppGpp crystallized with *B. anthracis* Hpt-1. Interaction diagrams generated by LigPlot for each ppGpp molecule in the asymmetric unit. Interactions for molecule A shown in (A) and interactions for molecule B shown in (B).

**Figure 2 - figure supplement 6.**
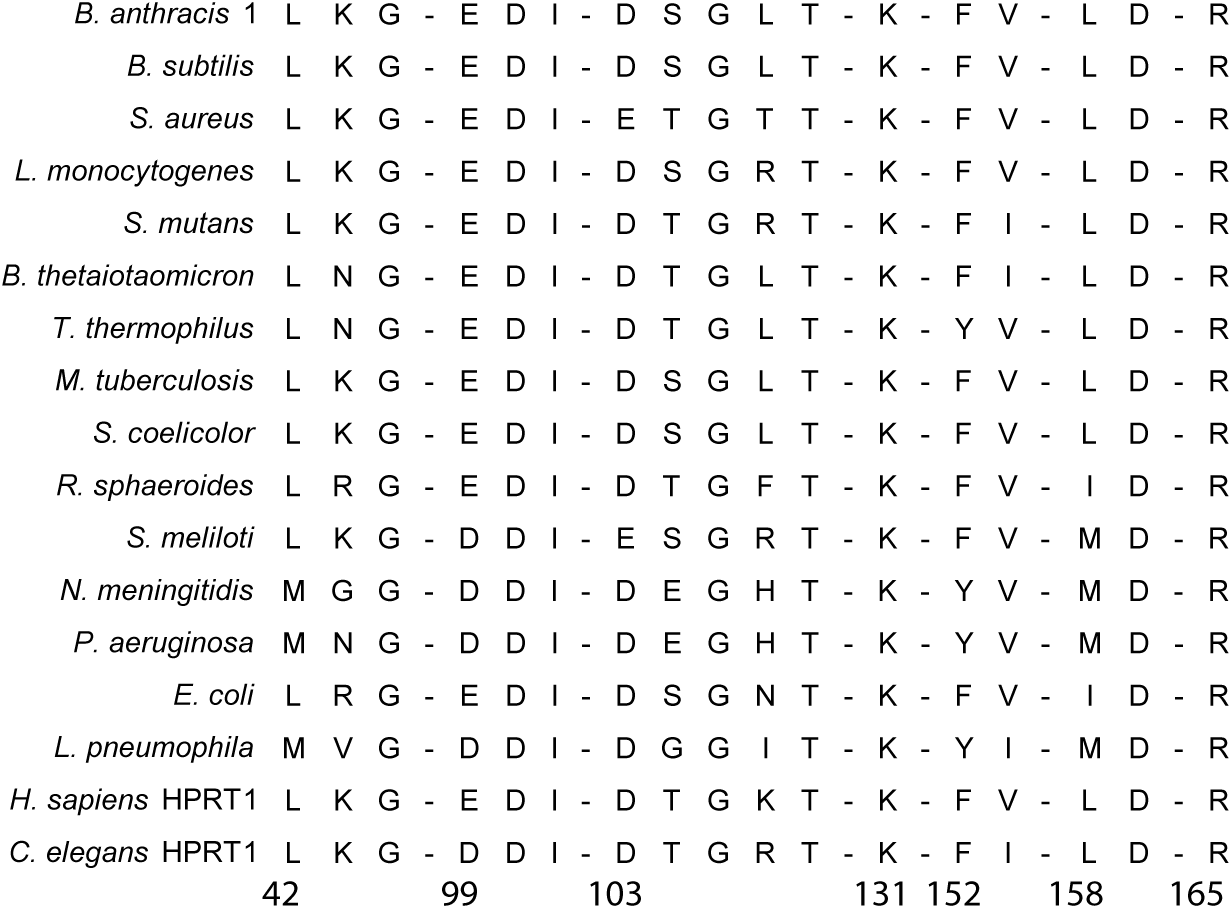
Conservation of ppGpp binding site across select bacteria and eukaryotes. (p)ppGpp binding residues are conserved. Alignment of (p)ppGpp binding residues from representative bacterial and eukaryotic HPRTs. The bacteria are a subset of the 99 HPRTs used to make the frequency logo in Figure 2F. Numbering is according to *B. anthracis* Hpt-1.

**Figure 3 - figure supplement 1.**
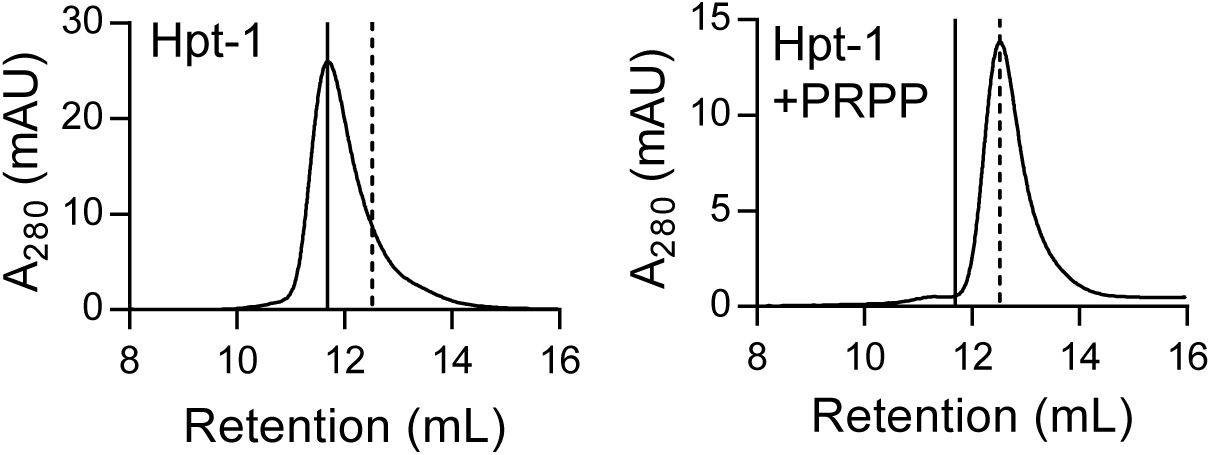
*B. anthracis* Hpt-1 is a tetramer without ligands and a dimer with PRPP. Size-exclusion chromatography of *B. anthracis* Hpt-1 without ligands (left) and with 1 mM PRPP in the mobile phase (right). The solid vertical line represents a tetramer, and the dotted vertical line represents a dimer.

**Figure 3 - figure supplement 2.**
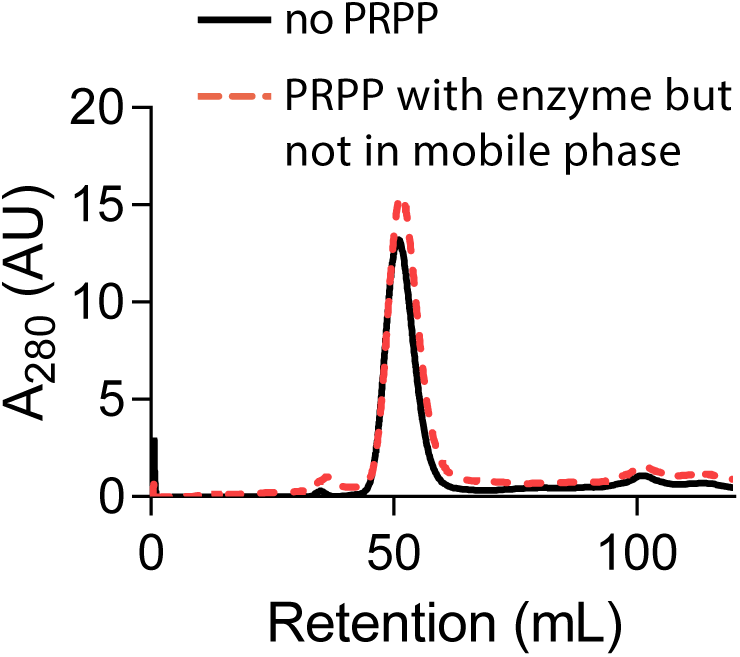
PRPP does not cause HPRT dimerization when not in the mobile phase. Size-exclusion chromatography of *B. subtilis* HPRT without PRPP (black solid line) or with PRPP in the enzyme sample but not in the mobile phase (red dotted line). Chromatography was performed with a HiPrep Sephacryl S-100 HR column.

**Figure 3 - figure supplement 3.**
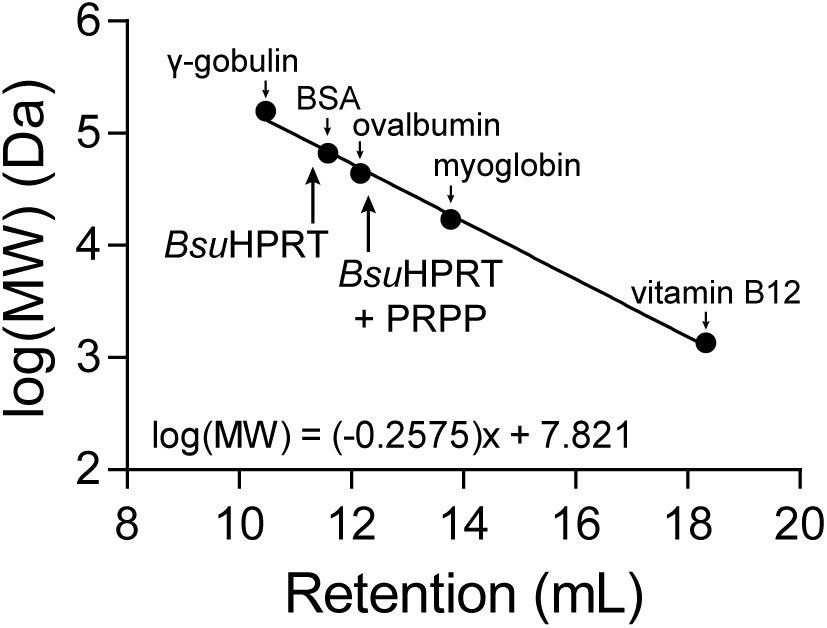
Molecular weight standards for size-exclusion chromatography. Molecular weight standards for the Superose 12 10/300 GL size exclusion column. The standards are: γ-gobulin (158 kDa), bovine serum albumin (BSA) (66.5 kDa), ovalbumin (44 kDa), myoglobin (17 kDa), and vitamin B12 (1.35 kDa). The equation is a linear regression of the log molecular weight in Daltons versus the retention volume of each marker. Arrows point to the retention volumes of *B. subtilis* HPRT with and without PRPP.

**Figure 4 - figure supplement 1.**
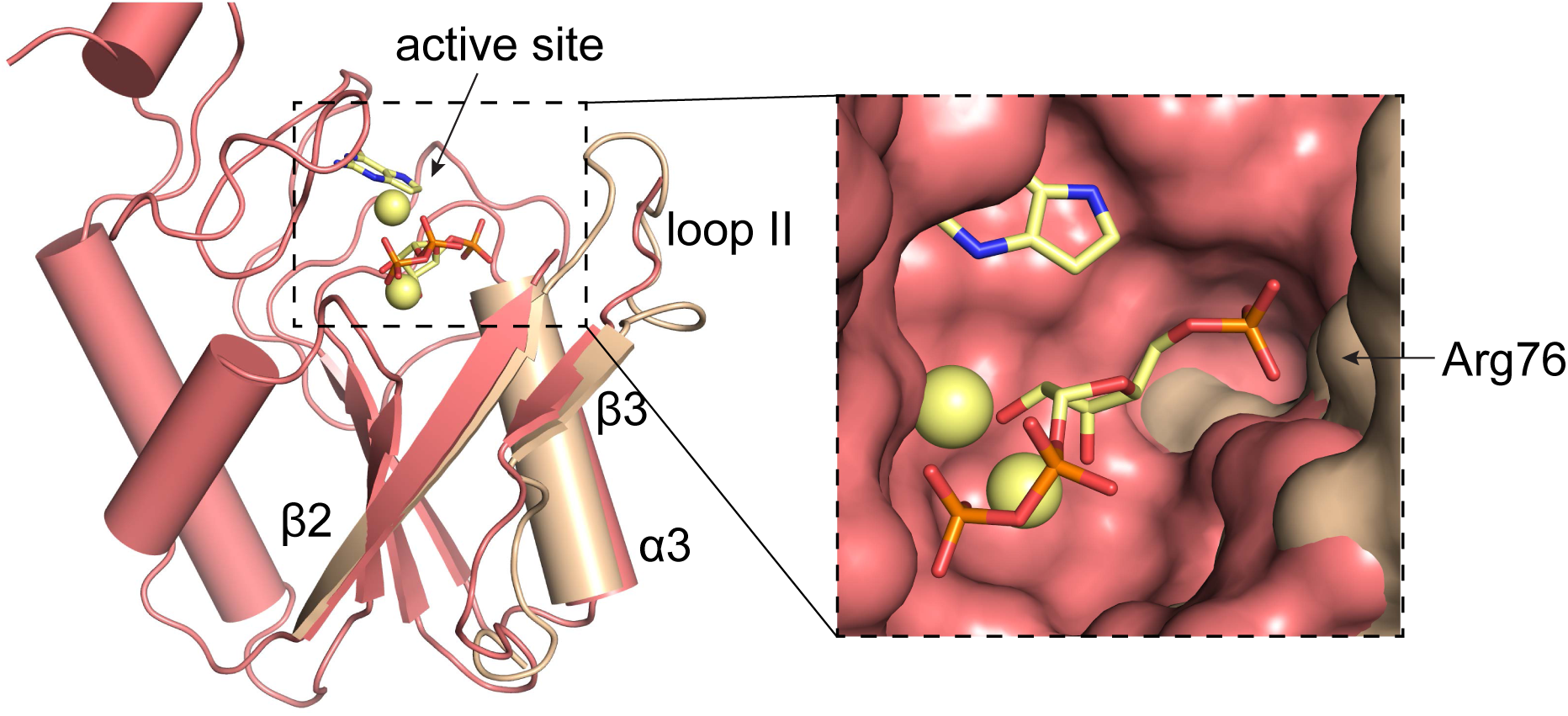
Overlay of *L. pneumophila* HPRT and substrates-bound *B. anthracis* Hpt-1. Overlay of substrate-bound *B. anthracis* Hpt-1 (salmon) and loop II, β3, and α3 from apo *L. pneumophila* HPRT (wheat; PDB ID 5ESW). PRPP and 9-deazaguanine are shown as yellow sticks and Mg^2+^ as yellow spheres. Inset shows surface view of the binding pocket with the *L. pneumophila* HPRT interface components.

**Figure 4 - figure supplement 2.**
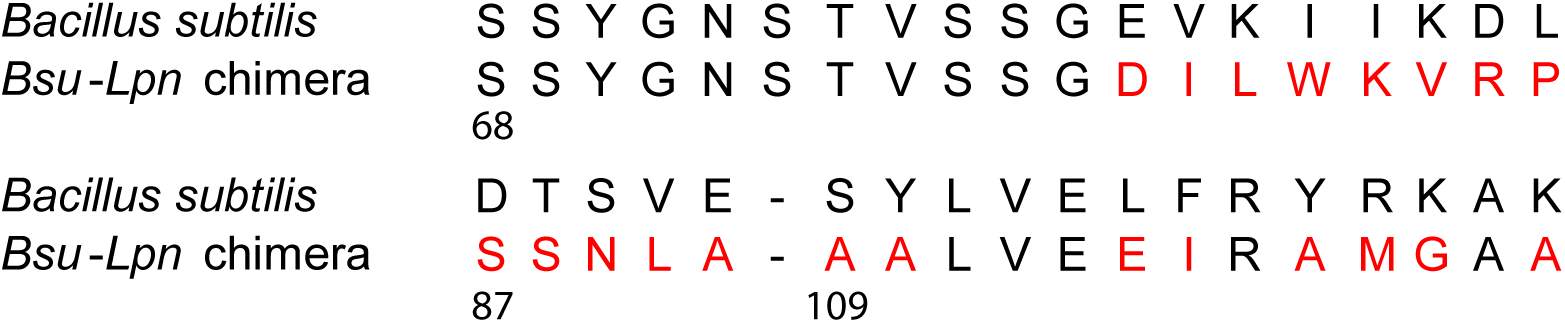
Residues replaced in *B. subtilis* HPRT for the *Bsu-Lpn* chimera. Residues at the *B. subtilis* (*Bsu*) HPRT dimer-dimer interface replaced with corresponding residues from *L. pneumophila* (*Lpn*) HPRT (shown in red).

**Figure 4 - figure supplement 3.**
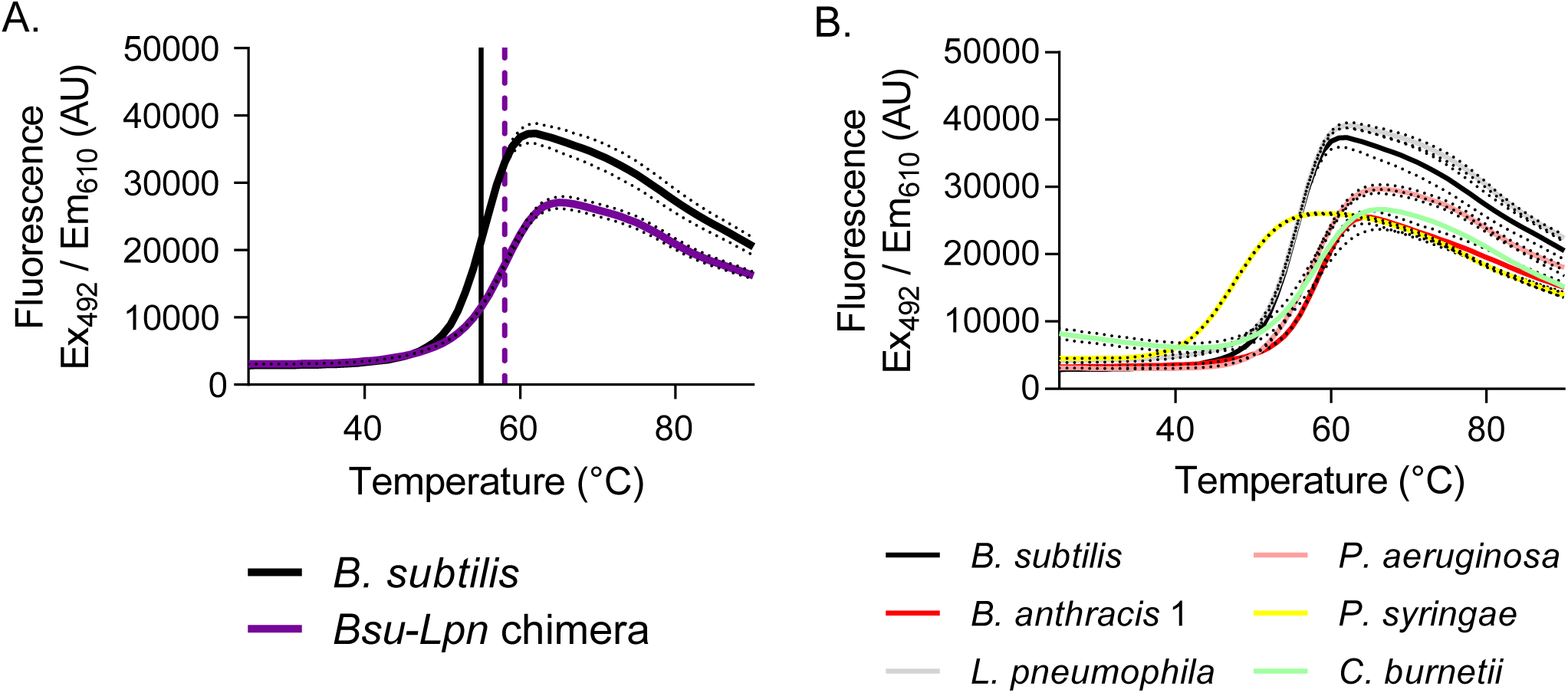
Stability of dimeric HPRTs is not compromised. **A)** Fluorescence intensity versus temperature of *B. subtilis* HPRT (black) and the *Bsu-Lpn* chimera (purple) in the presence of SYPRO orange. The vertical lines mark the melting temperature, determined from the first derivative of the melting curve (black solid line for *B. subtilis* HPRT and purple dashed line for the *Bsu-Lpn* chimera). Dashed lines represent SEM for reactions performed in triplicate. **B)** Fluorescence intensity versus temperature of bacterial HPRTs in the presence of SYPRO orange. Increased melting temperature indicates increased stability, and protein stability is independent of oligomeric state among the HPRT homologs tested. Dashed lines represent SEM for reactions performed in triplicate. *B. subtilis* HPRT data are the same as in panel A.

**Figure 5 - figure supplement 1.**
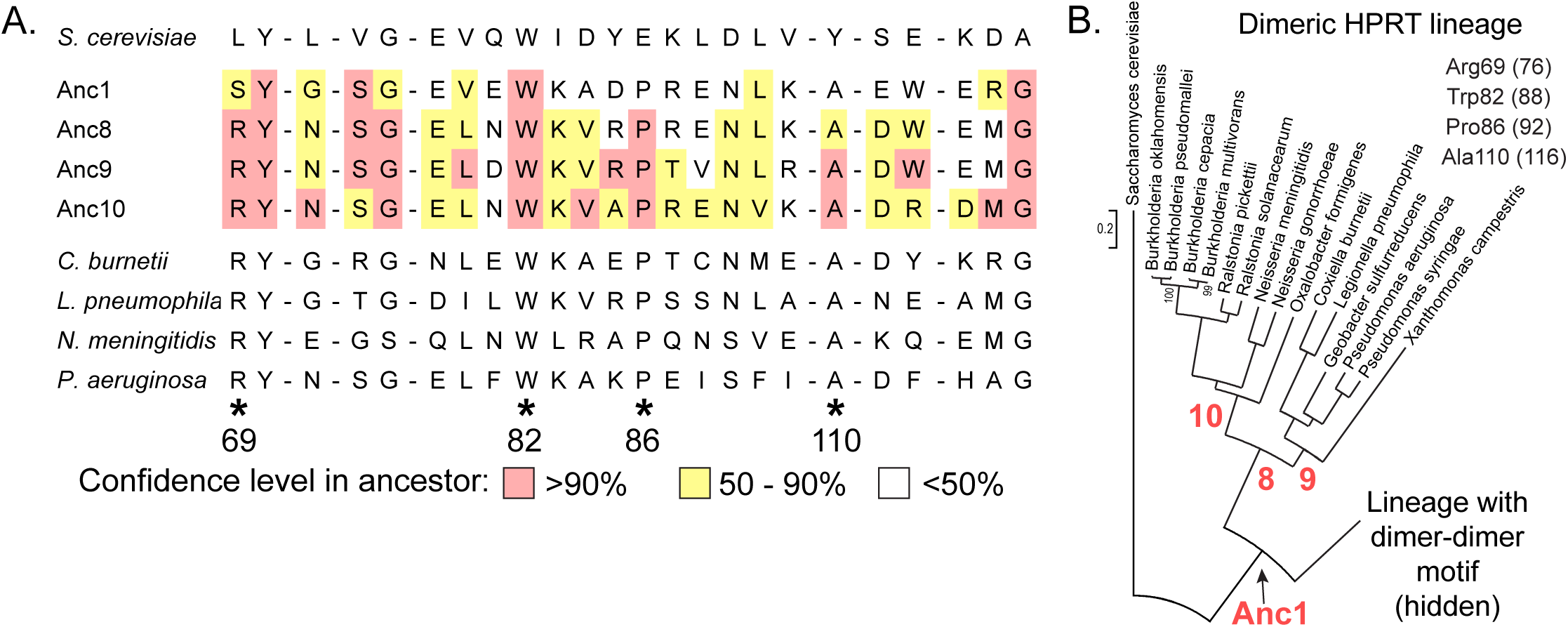
Evolution of dimer-dimer interface in the dimeric HPRT lineage. **A)** Alignment of putative dimer-dimer interface residues in dimeric HPRTs. Also shown are the residues in four ancestors to the dimeric HPRTs (Anc1, Anc8, Anc9, and Anc10). Note the conservation of Arg69, Trp82, Pro86, and Ala110 in dimeric HPRTs. For the ancestors, red indicates a high confidence level in the residue identity (>90%), yellow indicates a moderate confidence level (50-90%) and no color indicates a low confidence level (<50%). Numbering is according to *B. anthracis* Hpt-1. **B)** Lineage of HPRTs lacking the dimer-dimer interface motif. The tree is from Figure 5A with the remainder of the tree hidden. Instead of a Lys81-associated dimer-dimer interaction motif, the dimeric HPRTs share Arg69, Trp82, Pro86, and Ala110 residues at the interface. The numbers in parentheses refer to residue numbering in *L. pneumophila* HPRT. Red numbers indicate ancestral HPRTs (Anc1, Anc8, Anc9, and Anc10).

**Figure 5 - figure supplement 2.**
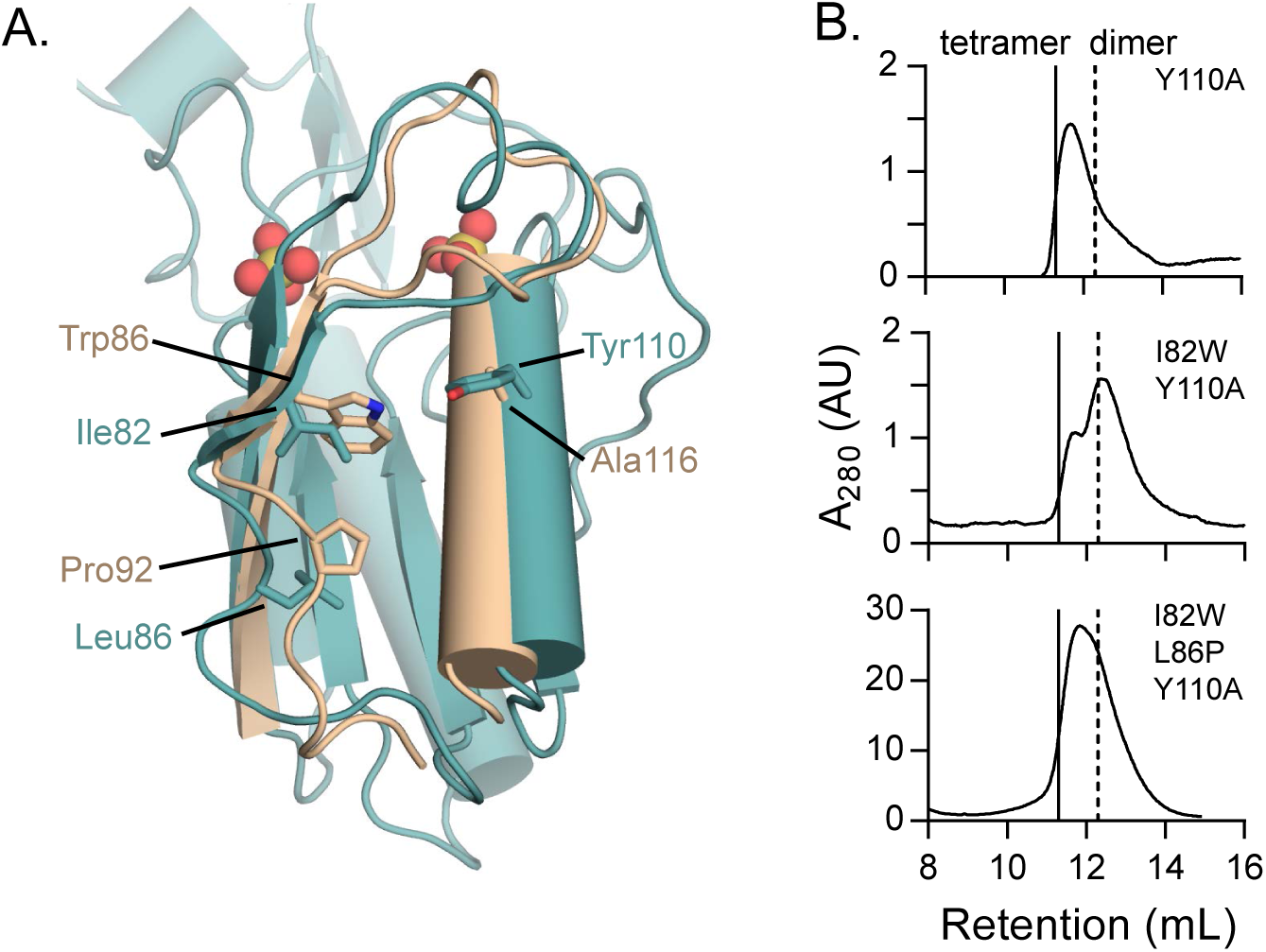
Role of residues conserved in the dimer-dimer interface of dimeric HPRTs. **A)** The role of interface residues conserved in dimeric HPRTs. Overlay of *B. anthracis* Hpt-1 (teal) and *L. pneumophila* HPRT (wheat; PDB ID 5ESW). Ile82, Leu86, and Tyr110 are shown from *B. anthracis* Hpt-1, and the homologous residues Trp88, Pro92, and Ala116 are shown from *L. pneumophila* HPRT. Trp88 packs between loop II and α3. Ala116, which replaces Tyr110, may help accommodate Trp88. Pro86 introduces a kink below loop II that may promote the packing of Trp88 against α3. **B)** Variants conserved in dimeric HPRTs weaken tetramerization. Size-exclusion chromatography of Y110A, I82W/Y110A, and I82W/L86P/Y110A *B. subtilis* HPRT variants. Solid vertical lines represent an expected HPRT tetramer and dotted vertical line represents an expected dimer.

**Figure 5 - figure supplement 3.**
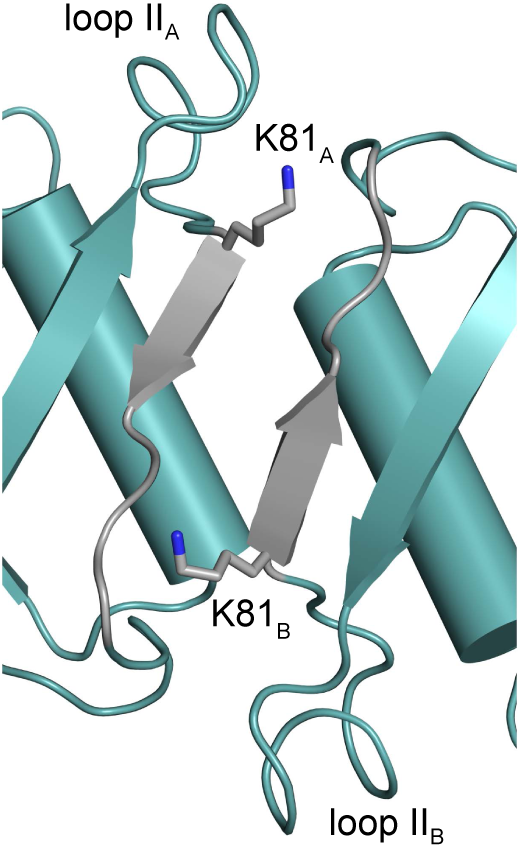
The conserved Lys81 bridges the dimer-dimer interface of tetrameric HPRTs. The structure of the dimer-dimer interface of *B. anthracis* Hpt-1 with the side chain of Lys81 shown as sticks. Regions colored gray correspond with residues 81-87 (see Figure 5B). The subscripts A and B refer to the subunit contributing the residues to the interface interaction.

**Figure 6 - figure supplement 1.**
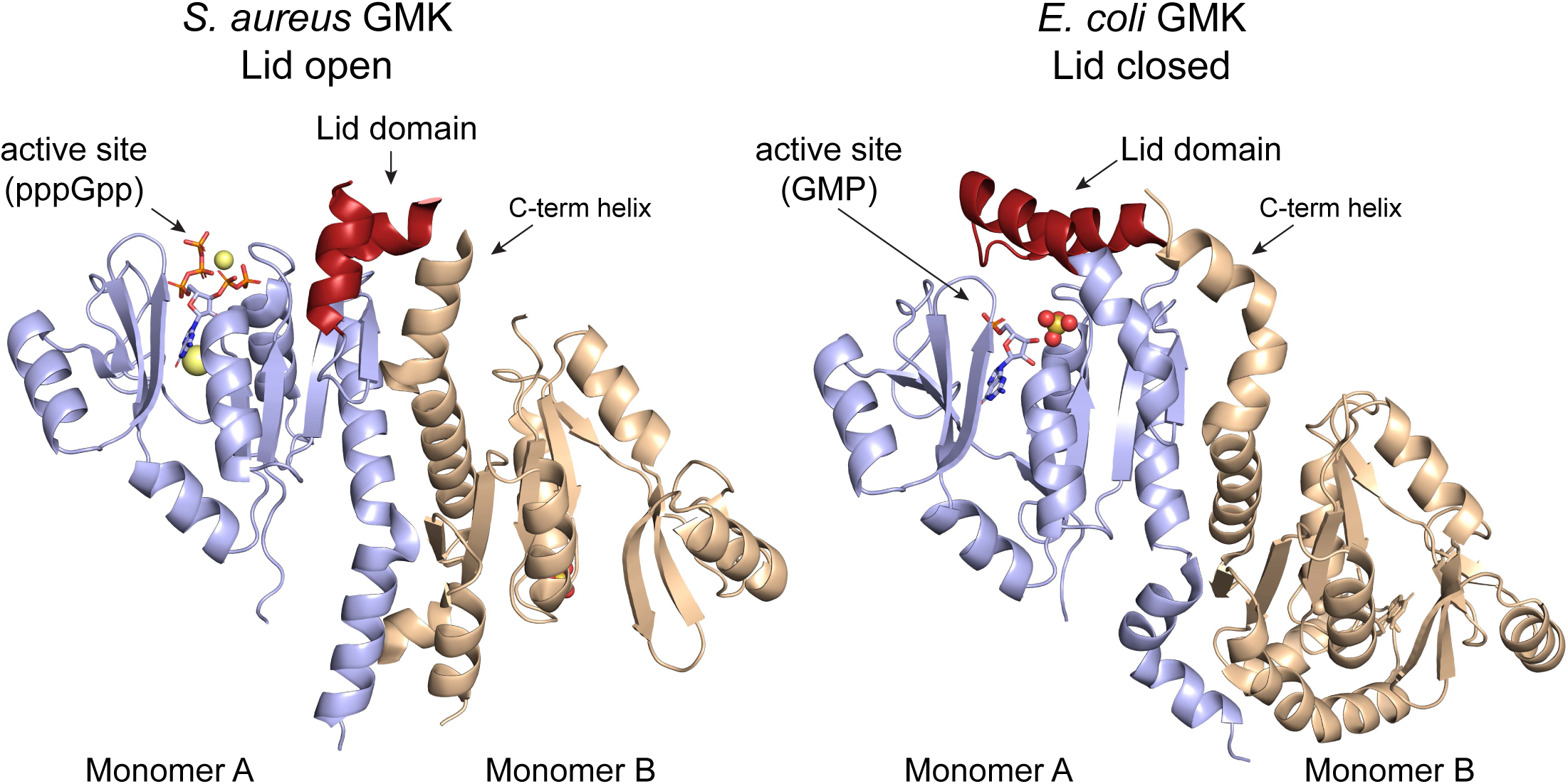
Role of oligomerization in altering the ligand binding site of guanylate kinase. The conformation of the (p)ppGpp binding site in guanylate kinase (GMK) may be related to its oligomerization. In *S. aureus* GMK (left; PDB ID 4QRH), an open lid domain (red) allows (p)ppGpp binding at the active site. However, a C-terminal helix in the second monomer of a GMK dimer interacts with the open lid domain. Nineteen residues in the lid domain and six residues in the C-terminal helix are unresolved. In *E. coli* GMK (right; PDB ID 2ANB), the closed lid domain occludes (p)ppGpp binding at the active site (bound to GMP). The *E. coli* C-terminal helix takes a different conformation that may help stabilize the closed lid domain.

## Source Data Files

Table 2 – Source Data 1. Validation report for PDB ID 6D9Q.

Table 2 – Source Data 2. Validation report for PDB ID 6D9R.

Table 2 – Source Data 3. Validation report for PDB ID 6D9S.

## Supplementary files

Supplementary file 1. Primers, plasmids, and strains.

